# Revision: Single-nanoparticle electrophoretic mobility determination and trapping using active feedback 3D tracking

**DOI:** 10.1101/2024.07.08.602591

**Authors:** Alexis Johnson, Kevin Welsher

## Abstract

Nanoparticles (NP) are versatile materials with widespread applications across medicine and engineering. Despite rapid incorporation into drug delivery, therapeutics, and many more areas of research and development, there is a lack of robust characterization methods. Light scattering techniques such as dynamic light scattering (DLS) and electrophoretic light scattering (ELS) use an ensemble-averaged approach to the characterization of nanoparticle size and electrophoretic mobility (EPM), leading to inaccuracies when applied to polydisperse or heterogeneous populations. To address this lack of single-nanoparticle characterization, this work applies 3D Single-Molecule Active Real-time Tracking (3D-SMART) to simultaneously determine NP size and EPM on a per-particle basis. Single-nanoparticle EPM is determined by using active feedback to “lock on” to a single particle and apply an oscillating electric field along one axis. A maximum likelihood approach is applied to extract the single-particle EPM from the oscillating nanoparticle position along the field-actuated axis, while mean squared displacement is used along the non-actuated axes to determine size. Unfunctionalized and carboxyl-functionalized polystyrene NPs are found to have unique EPM based on their individual size and surface characteristics, and it is demonstrated that single-nanoparticle EPM is a more precise tool for distinguishing unique NP preparations than diffusion alone, able to determine the charge number of individual NPs to an uncertainty of less than 30. This method also explored individual nanoparticle EPM in various ionic strengths (0.25-5 mM) and found decreased EPM as a function of increasing ionic strength, in agreement with results determined via bulk characterization methods. Finally, it is demonstrated that the electric field can be manipulated in real time in response to particle position, resulting in one-dimensional electrokinetic trapping. Critically, this new single-nanoparticle EPM determination and trapping method does not require microfluidics, opening the possibility for the exploration of single-nanoparticle EPM in live tissue and more comprehensive characterization of nanoparticles in biologically relevant environments.

## Introduction

Nanoparticles (NPs) are important materials being incorporated into a wide range of scientific and engineering applications, such as diagnostics, imaging, and drug delivery.^1, 2^ Due to their widespread application, detailed characterization of NPs in various contexts is critical. Despite extensive research on nanomaterials, few nanotechnologies reach clinical trials and rarely achieve commercial success in contributing to human health, with the notable exception of lipid nanoparticle-based vaccines developed in response to the SARS-Cov-2 pandemic.^3, 4^ Challenges faced by nanomedicine are often rooted in the lack of robust characterization in complex biological environments. The abundance of various proteins and salts in biological sera leads to changes in surface properties that may impact the transport of NPs to target sites, decrease cellular uptake of NPs, and even increase cytotoxicity and autophagic response.^4^ NP characterization methods that faithfully report changes to surface properties are critical for developing and understanding successful nanomedicines, particularly *in vivo*.

Light-scattering-based techniques have become the workhorses of nanomedicine development. Dynamic light scattering (DLS) is a bulk technique, using scattered light from diffusing objects in dilute solutions, aided by an autocorrelation function, to measure the hydrodynamic radius of nanoscale particles.This technique is robust for reporting the ensemble-averaged radius of particles in monodisperse solutions but faces many challenges with polydispersity.^5^ The scattering signal from large particles or small aggregates can easily eclipse single molecules.^6^ Biological solutions contain various proteins and other macromolecules that may induce NP aggregation, leading to a high degree of polydispersity and difficulties in assessing appropriate dosage, toxicological findings, and identifying therapeutic mechanisms of action.^7^

Measurement of size and aggregation state via DLS is only an indirect readout of the NP surface properties. Measurement of the electrophoretic mobility can reveal more detailed information about surface chemistry. Electrophoretic mobility (EPM) is the velocity of colloidal particles in an external electric field and is highly sensitive to NP size, surface characteristics, and solubility. EPM is often reported via the electrostatic surface potential or zeta potential (ZP) using Smoluchowski’s formula for rigid, spherical colloids.^8^ EPM is measured using electrophoretic light scattering (ELS), operating in the same fashion as DLS, but with the addition of an externally applied electric field. Brownian motion, or the “random walk” of particles in solution, is altered by the electric field to probe the surface properties of NPs. Both EPM and ZP are used to assess colloidal stability, protein corona formation, and protein-nanoparticle interactions. For example, Santander-Ortega and coworkers used EPM to assess the colloidal stability and adsorption of IgG antibodies on polymeric NPs at physiological pH to simulate a drug delivery system with targeting abilities.^9^ Malburet *et al*. elucidated the physiochemical parameters of polydisperse lipid nanoparticles (LNPs) used as a delivery system for mRNA vaccines by measuring EPM and using it to calculate ZP and surface charge density.^10^ Unfortunately, methods used to characterize EPM in biological contexts suffer from the same pitfalls as DLS. The EPM reported is a product of an ensemble average that is likely to be biased towards large particles in solution, similar to DLS. The advancement of nanomedicine requires new, robust, and unbiased methodology.

As an ensemble measurement, ELS struggles with accurately characterizing heterogenous solutions, where critical sub-populations can be obscured. Single-particle approaches are key to capturing these potentially critical sub-populations and dynamics. To address this shortcoming, single particle methods with the capability to measure EPM and ZP have emerged. For example, tunable resistive pulse sensing (TRPS) traps single particles in nanosized stretchable pores, providing information about size and EPM. Although TRPS is a commercially successful technique, it encounters complications including, easily clogged nanopores, requirement of calibration and standardization and unfavorable pore-particle interactions.^11, 12^Other approaches to the measurement of size and EPM at the single-particle level are microscopy-based methods developed by Choi et al.^13^ and Oorlynck et al (Table S1).^14^ These image-based particle tracking methods monitor particle behavior along one axis in response to an electric field to measure EPM in a home-built microfluidics chamber. Image-based methods often have a limited, predetermined observation volume, which is often a result of the limited axial range.^15^ This is apparent in these methods as the Z axial range for each technique is limited to 8 μm and 1.4 μm, respectively. Even in shallow microfluidics chamber, this can lead to variations in EPM measurements due to the axial position of the particle. Furthermore, these single-particle EPM techniques have a limited observation time. Many biological processes that EPM could potentially monitor happen on the order of minutes to hours.^15^ TrACE, presented by Choi *et al*., primarily follows single particles for 1 second at a time. The laser scanning microscopy method by Oorlynck and coworkers uses a similar observation time, combining power spectral densities of multiple particles for longer traces (i.e. 81 particles are combined to yield 30 s of data). Alternatively, the Anti-Brownian Electrokinetic trap (ABEL trap) uses electrokinetic feedback to prolong the observation of single particles with their microfluidic confined axial range (∼700 nm) and can subsequently determine the electrokinetic mobility of single particles.^16^ The ABEL trap can estimate the diffusion coefficient and electrokinetic mobility by trapping single particles but does so at very high field strengths. For future inquiry into how EPM and diffusion coefficient might change in complex environments (i.e. cells, tissue), the ABEL trap is not ideal because of the likelihood of electroporation.

While existing methods rely on confining particles to limited observation volumes or limited observation times, real-time 3D single-particle tracking (RT-3D-SPT) can lock onto individual particles as they diffuse over long 3D distances, collecting long trajectories of single NPs with millisecond or better temporal resolution. Importantly, this technique is easily translatable to complex biological settings. Here, we demonstrate that 3D Single-Molecule Active Real-Time Tracking (3D-SMART) is an ideal method for single-NP EPM determination.^17^ 3D-SMART locks onto fluorescent polystyrene nanoparticles (PS NPs) in solution as they diffuse using active-feedback tracking (Fig. 1). A dynamic observation volume follows the particle of interest as it diffuses and uses the signal from the single-photon counting detector, removing the temporal limitations imposed by a camera exposure and readout times in image-based tracking methods. Previously, 3D-SMART has been able to track viral contacts with the cell surface,^18^ characterize *in situ* protein corona formation,^19^ determine growth kinetics of single polymer particles in solution,^20^ and characterize LNP diffusion through mucus suspensions.^21^ This technique has proven capable of extracting precise nanoparticle characteristics in complex environments. With the addition of an applied electric field, 3D-SMART provides a 3D trajectory at a kHz sample rate, simultaneously revealing single particle parameters such as diffusion coefficient, hydrodynamic radius, and electrophoretic mobility, which is new to this work.

**Figure 1:**
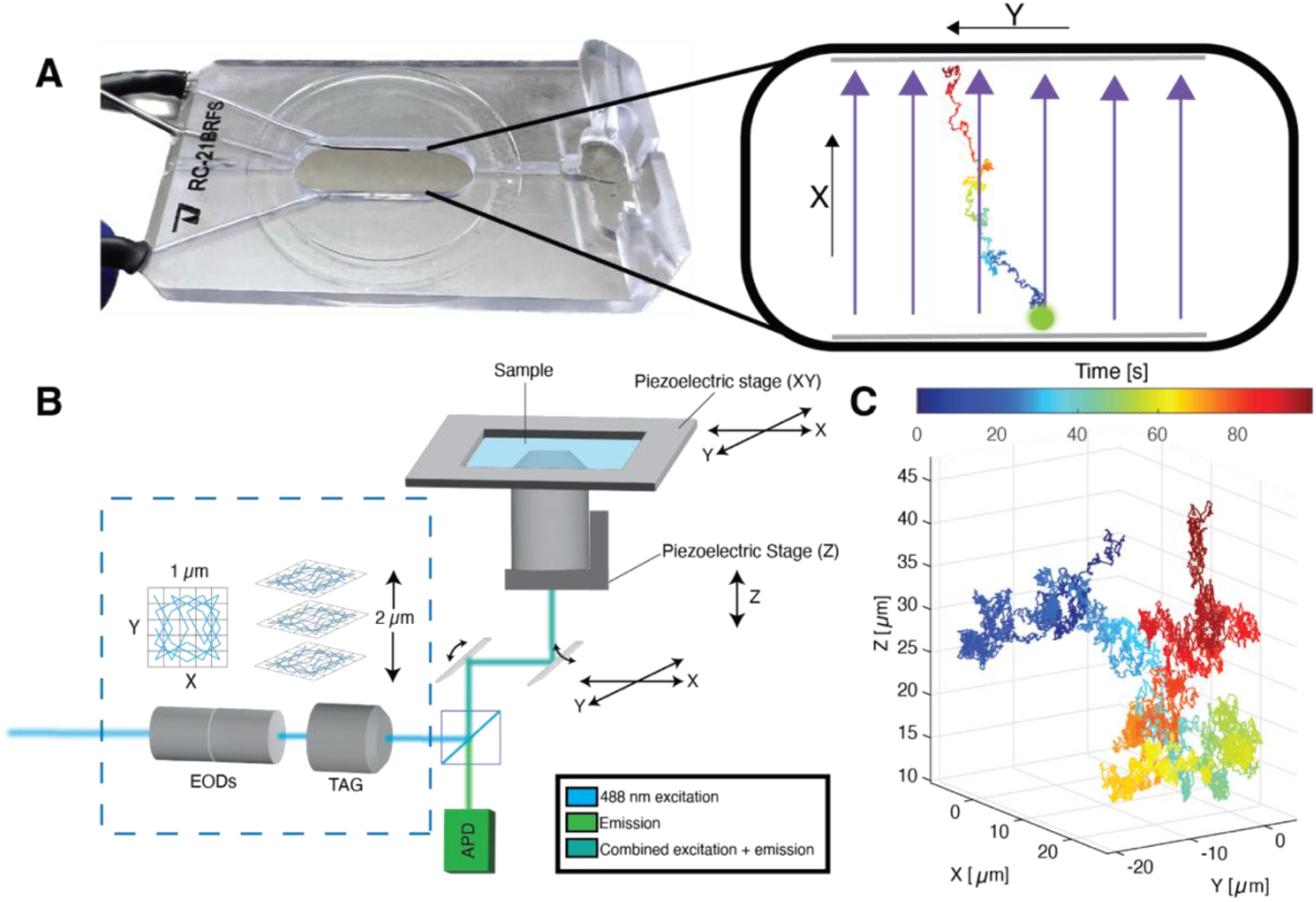
Experimental scheme for single-nanoparticle electrophoretic mobility measurements. (A) Field Stimulation open bath chamber (Warner Instruments RC-21BRFS, 18 × 6.3 × 2.3 mm). Inset: Cartoon scheme of single nanoparticle under electric field. (B) 3D-SMART microscope illustration. Detailed included in Methods section. (C) Example 3D trajectory of free diffusing nanoparticle in water, captured by 3D-SMART.

## Results and Discussion

### Active-Feedback 3D Single-Particle Tracking with Electric Field Manipulation

3D-SMART achieves real-time tracking of single fluorescent nanoparticles through rapid position estimation within a 3D observation volume, which is then used as the input to a feedback loop that recenters the particle in the observation volume. The 3D volume is created by the electro-optical deflectors (EODs) to deflect a 488 nm laser in X and Y in a knights tour pattern^22^ at 20 μsec per spot and 500 μsec for the entire 5 × 5 grid. Along the Z axis, the focus is deflected in a sine wave at ∼70 kHz by a tunable acoustic gradient (TAG) lens. Together, the EODs and TAG lens create a rapid scan over a 1 × 1 × 2 μm scanning volume. Position estimates are made using photon arrival times recorded by the avalanche photodiode (APD) and a Kalman filter implemented on a field programmable gate array (FPGA, NI PCIe-7852R). The position estimate is fed into an integral feedback controller to center the observation volume on the diffusing nanoparticle via piezoelectric nanopositioners or galvo scanning mirrors. In addition to the standard configuration of 3D-SMART, measurements of electrophoretic mobility require an applied electric field. Here, an electric field is applied using the analog output of the FPGA and is controlled via National Instruments LabVIEW software. For 3D-SMART, experimental conditions include a small sample volume (∼300 μL) in an open bath chamber with parallel platinum electrodes, spaced 6.3 mm apart, capable of field stimulation (Warner Instruments RC-21BRFS, Fig. 1A). This chamber houses freely diffusing NPs in solution for the duration of the experiment under various field stimulation modes, allowing Brownian motion for PS NPs in water to be recorded in real-time by 3D-SMART (Fig. S1). The 18 × 6.3 × 2.3 mm chamber makes is unlikely for NPs in the dilute solution to encounter the chamber walls in X and Y. Any alternations in diffusive motion or electrophoretic mobility due to contact with the side walls in two-dimensions are assumed to be negligible. For Z, the centered tracking position is set to 20 microns above the coverslip. Occasionally particles encounter the glass cover slide (precipitating on the surface or momentary contacts) but these events are apparent in the 3D trajectories and these trajectories can easily be excluded.

The maximum applied voltage across the sample for the majority of the experiments below is 2 V, corresponding to a field strength of 3.2 V/cm. The voltage was set after testing the response of the 3D-SMART system at different voltages ranging from 1 to 10 V, measuring the resulting drift velocity, and comparing it to the expected outcome (Fig. S2). For 200 nm and 100 nm particles, lower voltages yielded a more accurate drift velocity. Nevertheless, the experimental drift velocities are within one standard deviation of the expected drift velocities for majority of the voltage range (Figure S2A and B). The deviations are likely due to position estimation inaccuracies at high velocities.

Figure 2A-C demonstrates the modes of voltage application utilized in the current study: zero applied voltage corresponding to Brownian motion (0 V, Fig 2A), constant voltage (+2 V Fig. 2B), and oscillating voltage between -2 V and + 2 V at a frequency of 1 Hz (±2 V, Fig. 2C). In the zero-field condition, Brownian motion is observed, as shown for a 194 nm polystyrene (PS) NP diffusing for approximately 90 seconds in Figure 2D. The 3D trajectories (Fig. 2D-F), which are extracted from the readout of the piezoelectric nanopositioners and galvo mirrors, show the movement of the particle overall. The X-position (purple), which corresponds to the axis of field stimulation overlaid in green, is isolated from the trajectory (Fig. 2G-I) for mean squared displacement (MSD) analysis (Fig. 2J-L) to illustrate the effect of the field actuation on particle motion.

**Figure 2:**
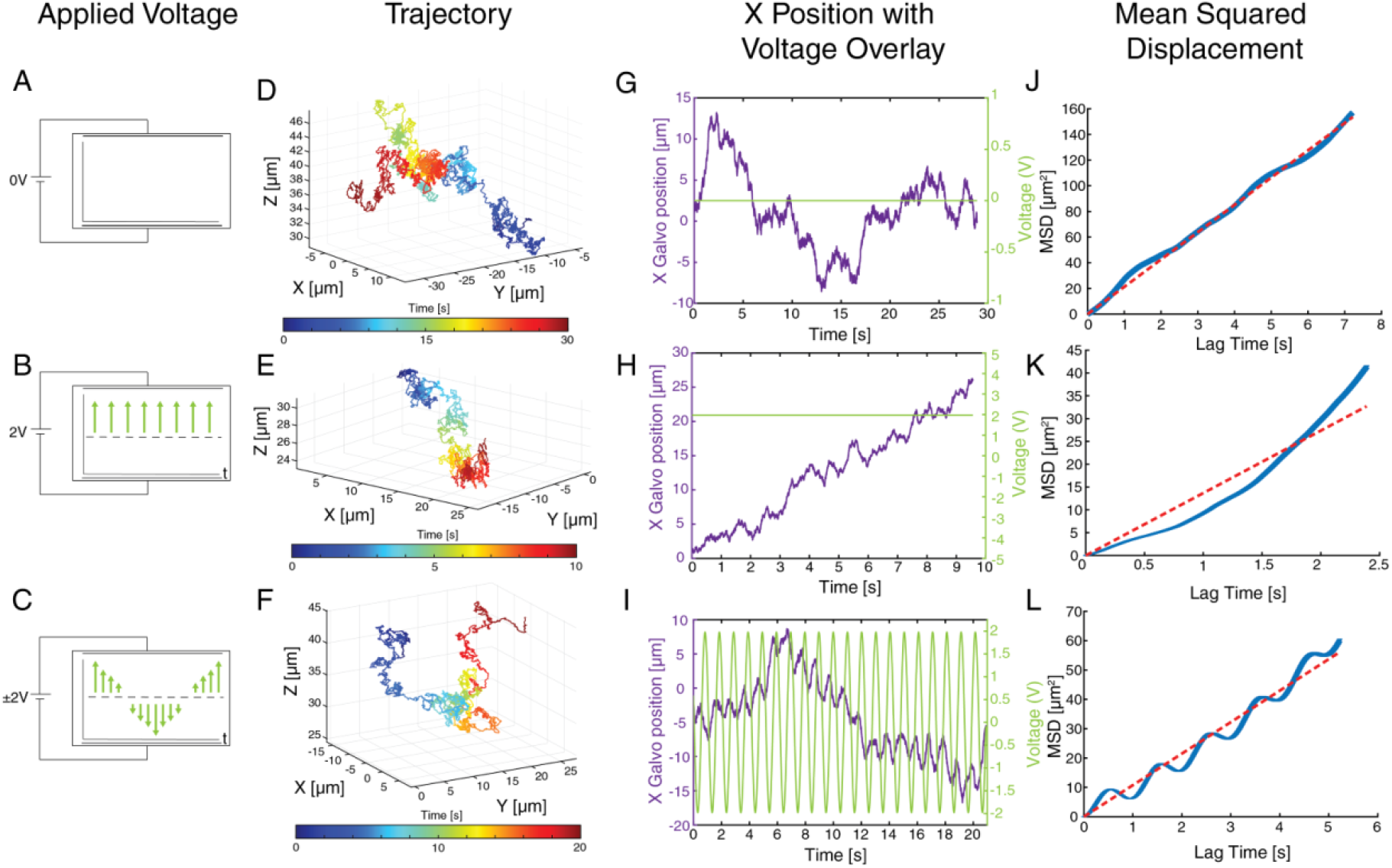
Three-dimensional motion of nanoparticles in solution under various applied voltage conditions. (A-C) Illustration depicting the voltage conditions applied along the X direction: 0 V, constant application of +2 V, and sinusoidal oscillation between +2 V and –2V at a frequency of 1 Hz. (D-F) Representative 3D trajectories of 194 nm unfunctionalized polystyrene nanoparticles diffusing under the field conditions specified in (A-C). (G-I) X position of nanoparticles versus time (purple) overlaid with the field conditions specified by (A-C) versus time (green). (J-L) Mean squared displacement versus lag time of particles under the field conditions specified in (A-C). The dotted line denotes the expected linear trend of a freely diffusing particle.

When applying an electric field along the X-axis, the behavior of a 194 nm PS NP undergoes noticeable changes along the manipulated axis. The data above clearly establishes that the expected Brownian motion is perturbed in response to the applied electric field. Upon examining the overlay of the applied voltage and the position of the particle along the field axis, it can be observed that the particle moves in direct response to the field. In contrast to the zero-field condition which is freely diffusing (Fig. 2G), a constant voltage of +2 V along the X direction of the sample chamber “pushes” the particle in one direction correlating with the applied electric field (Fig. 2H). The applied field results in a deviation from Brownian motion, which is clearly observed in the non-linear slope of the MSD plot (Fig. 2K). The MSD curve slopes upwards with increased lag time, taking on a convex shape. This shape is characteristic of particle drift or directed motion, which strongly suggests that the particle is drifting due to the constant applied voltage. The relationship between the applied field and the particle motion can be seen most clearly in the oscillating field case (Fig. 2I). A tight correlation is seen between the 1 Hz applied field and the particle position. Higher oscillation frequency can also be easily readout from the particle position data but does not have a significant effect on EPM determination so 1 Hz was used across all of the following experiments in this work (Figure S3). This is further confirmed using a Welch’s power spectral density estimate performed on the X position data as compared to the non-actuated Y axis (Fig. S4). The MSD takes on an unusual shape that is not one of the typical behaviors used to classify the motion of molecules in solution (Brownian, directed or confined motion). The MSD versus lag time takes on the same oscillating behavior as the applied field. The presented data makes it apparent that 3D-SMART can track the manipulated position of NPs in solution in response to an electric field. The response of the particle position due to the change in voltage is the basis for measuring electrophoretic mobility. The EPM is determined using a maximum likelihood estimation (Equation 9) which utilizes particle information extracted using 3D-SMART (particle displacement, drift velocity, etc.) and electric field strength calculated from the applied voltage monitored by the FPGA.

MSD plots for the non-actuated axes (Y and Z, Fig. S5) are used to accurately determine the diffusion coefficient and subsequently the hydrodynamic radius via the Stoke-Einstein relation (Equation 1 and 2). 3D-SMART allows for measurement of EPM while monitoring the aggregation state of single nanoparticles. Further details of how EPM is calculated is in the Methods section below.

### Precision of electrophoretic mobility measurement scales with trajectory duration

After collecting trajectories under the influence of an applied field and calculating the underlying EPM, the data were next evaluated to determine the factors related to the precision of the EPM extracted parameters. It was observed that the trajectory duration is the critical factor in EPM precision. To reach a relative uncertainty of ∼10%, single-particles trajectories must exceed 20 seconds in tracking duration when applying 2 V. Trajectories collected by 3D-SMART are occasionally short (<20 seconds) because of particle dimness, diffusion speeds beyond microscope tracking capabilities, or reaching a physical limit (encountering the edge of the stage or galvo range). As the cumulative tracking duration increases, the relative uncertainty of the calculated EPM decreases for 200 nm NPs, hitting ∼10% at 20 seconds and ultimately reaching 4% at 180 seconds (Fig. 3A). The same trend is observed for 100 nm NPs (Fig. S6). Given this requirement for trajectory duration, the oscillating field approach is preferred over the constant field approach. Under constant field, particles will quickly drift across the active tracking range, reducing the observation time. The oscillating field gives no net bias on the particle’s position, enabling longer observation. The majority of trajectories in an oscillating field (93 out of 96) meet this cut-off (Fig. 3B). Trajectories shorter than 20 seconds were excluded from further analysis, and the average relative uncertainty of retained oscillating field trajectories at 20 seconds is 11%.

**Figure 3:**
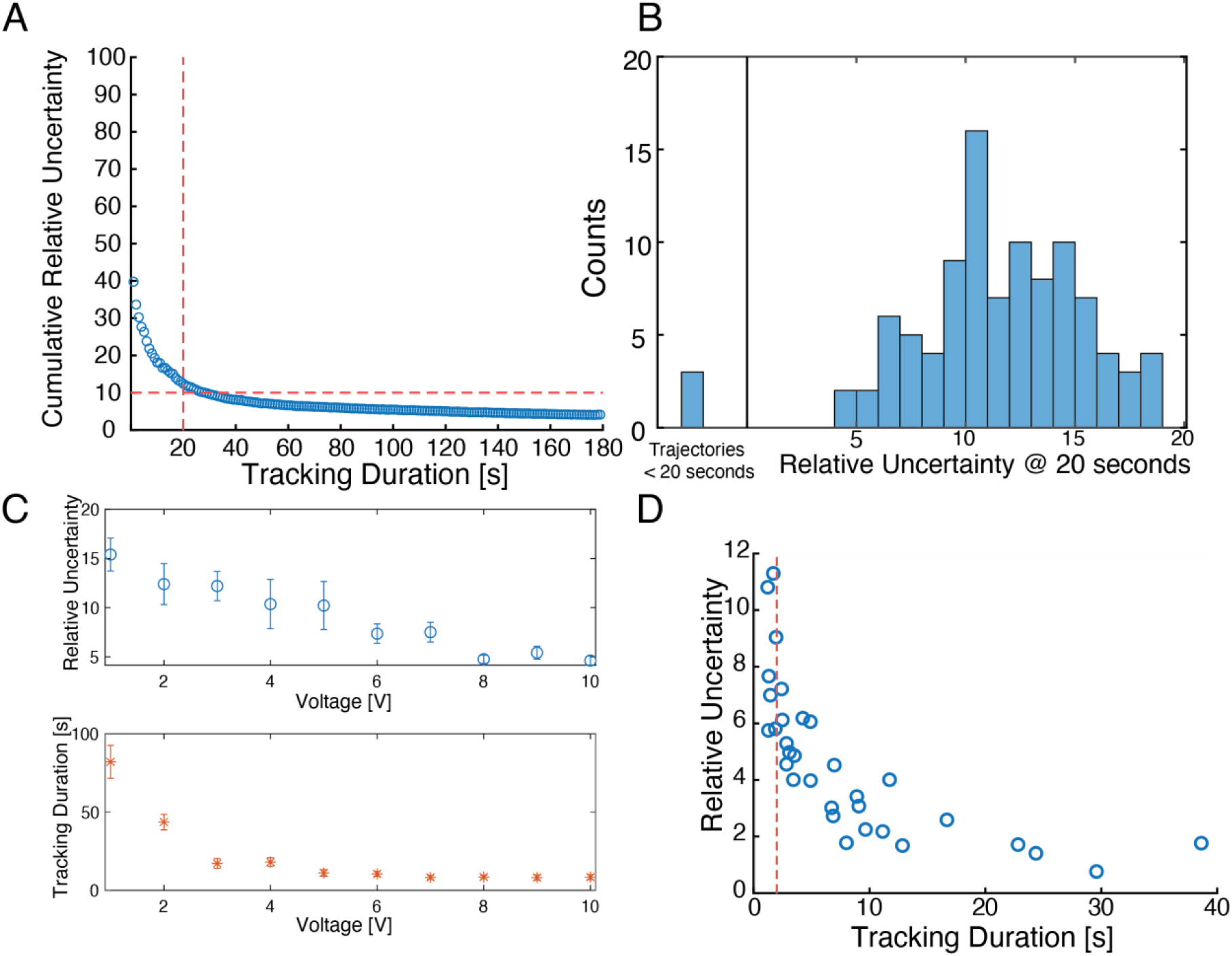
Relative EPM uncertainty versus trajectory duration. (A) Relative uncertainty of electrophoretic mobility for 196 nm carboxyl-functionalized polystyrene nanoparticles calculated at each second of continuous tracking. The dotted red line denotes the desired cutoff criteria: 20 seconds of single-nanoparticle tracking (vertical) and 10% relative measurement uncertainty (horizontal). (B) Histogram of relative uncertainty values reached for individual trajectories at 20 seconds of tracking (N = 96). Precision of voltage range. (C) Top: Relative uncertainty (95% confidence interval) of 198 nm COOH PS NPs in 1-10 V voltage range. With error bars representing standard error. Bottom: tracking duration of 198 nm COOH PS NPs in voltage range 1-10 V. With error bars representing standard error. (D) Relative uncertainty vs tracking duration of 198 nm COOH PS NPs at 10 V. Dashed red line denotes 2 seconds of tracking.

In cases of rapid data collection and limited tracking duration, this 20-second cut off can be avoided by using a higher applied voltage (Figure S7A). Higher precision can be achieved at higher voltages, but it sacrifices tracking duration and limits long-term observations (Figure 3C). When averaging the relative uncertainties and tracking durations of 30 trajectories, each collected at voltage 1-10 V, both parameters decrease with increasing voltage. When considering all trajectories at 10 V, 200 nm particles have a mean tracking duration of 8 seconds with 5% precision (N = 32), and 100 nm particles have a mean tracking duration of 4 seconds with a 9% precision (N = 36, Figure S7A). If only considering trajectories less than 2 seconds at 10 V, 3D-SMART reaches conditions comparable to previous methods with better precision. 3D-SMART reaches 11% for 100 nm particles (Figure S7B) and 8% for 200 nm particles (Figure 3D).Previous single molecule methods for EPM determination have achieved 5% for 500 nm nanoparticles^14^ and 15% precision for 100 nm nanoparticles^13^ as calculated from the reported standard error converted to 95% confidence interval (relative uncertainty) (Table S1).

Across the voltage range, EPM increases by ∼0.4×10^-8^ m^2^V^-1^s^-1^, approximately one standard deviation of the mean for 198 nm particles (Figure S3C). For 102 nm particles this change is ∼0.6×10^-8^ m^2^V^-1^s^-1^, also approximately one standard deviation of the mean (Figure S2D). This most likely arises from the inaccuracies in drift velocity at higher voltages and is amplified when converted to electrophoretic mobility (Equation 5).

### Single nanoparticle determination of electrophoretic mobility

Electrophoretic mobility is affected by a particle’s size, shape and charge. In this work, active-feedback single-nanoparticle EPM was applied to two different types of NPs: unfunctionalized polystyrene (PS) and carboxyl functionalized polystyrene (COOH-PS) NPs to explore how EPM is affected by size and charge at a particle-by-particle basis. The nominal diameter of the unfunctionalized PS NP used is 194 nm, as reported by the manufacturer (Bangs Laboratories) via DLS. The reported diffusion coefficient by 3D-SMART as measured by using the MSD of the non-actuated axes (Y and Z), is 2.45 ± 0.96 μm^2^/s (diameter = 202 ± 82). The diffusion coefficient determined by our experimental DLS measurement is 2.56 ± 0.8 μm^2^/s (Table 1). More replicates of the diffusion coefficient of 194 nm unfunctionalized PS NPs for 3D-SMART and DLS are included in Tables S3 and S10 and Figure S8.

**Table 1:** Electrophoretic mobility and diffusion coefficient results from 3D-SMART and ELS. Mean ± s.d.

| Particle | 3D-SMART |  | ELS |  |
| --- | --- | --- | --- | --- |
| | EPM ($\times 10^{-8} \text{ m}^2\text{V}^{-1}\text{s}^{-1}$) | Diffusion Coefficient ($\mu\text{m}^2\text{s}^{-1}$) | EPM ($\times 10^{-8} \text{ m}^2\text{V}^{-1}\text{s}^{-1}$) | Diffusion Coefficient ($\mu\text{m}^2\text{s}^{-1}$) |
| **194 BARE** | $-3.85 \pm 0.2$ | $2.45 \pm 0.96$ | $-3.29 \pm 2.7$ | $2.56 \pm 0.8$ |
| **198 COOH** | $-4.08 \pm 0.3$ | $2.37 \pm 0.95$ | $-4.08 \pm 0.52$ | $2.55 \pm 0.8$ |
| **196 COOH** | $-3.91 \pm 0.6$ | $2.49 \pm 1.0$ | $-1.13 \pm 0.4$ | $2.19 \pm 0.3$ |
| **102 COOH** | $-3.5 \pm 0.4$ | $4.8 \pm 1.7$ | $-4.43 \pm 0.83$ | $4.68 \pm 2$ |

The nominal diameters of the COOH-PS NPs characterized in this work are 102 nm, 196 nm and 198 nm. We use subtle and large differences in diameter NPs to monitor the role that size plays in this method and the EPM of each population. The diffusion coefficients for 198 nm, 196 nm and 102 nm COOH-PS NPs are 2.37 ± 0.95 (diameter = 208 ± 76 nm, N = 92), 2.45 ± 1 (diameter = 202 ± 83 nm, N = 79) and 4.8 ± 1.7 (diameter = 102 ± 48 nm, N = 73) μm^2^/s, respectively. The distribution of these populations and addition replicates can be found in the SI (Fig S9-11). According to our experimental DLS measurements, the diffusion coefficients of these NPs are 2.55 ± 0.8 (diameter = 193 ± 82), 2.19 ± 0.3 (diameter = 199 ± 82) and 4.68 ± 2 (diameter = 104 ± 54) μm^2^/s. For the monodisperse samples, characterized within this body of work, DLS has high precision and accuracy but lacks the ability to correlate the size of each particle to other parameters such as EPM and particle intensity. This allows us to identify aggregates and outliers in each population. 3D-SMART highlights the distribution while simultaneously measuring diffusion coefficient and electrophoretic mobility.

As compared to other single-nanoparticle methods of simultaneously measuring size and electrophoretic mobility, 3D-SMART has superior accuracy for particle radius. 3D-SMART has an accuracy of ∼97% for the particle diameters reported above, as compared to 75%^13^ and 64%^14^ with other methods. These accuracies also include measured particles more than twice the size of the NPs evaluated by 3D-SMART. 3D-SMART reaches ∼40% precision in terms of RSD for the reported mean diameter but for majority of the populations stay within the range of ∼10% accuracy. This is a well-established method of single-nanoparticles size estimation, the addition of EPM adds an extra dimension for characterizing individual nanoparticles.

The electrophoretic mobility is determined by the actuated (X) axis of the 3D position data and the applied electric field using a maximum likelihood estimation (MLE, Equation 9). This is performed on each individual trajectory to pull out the single-NP EPM. For 194 nm PS NPs, the average EPM of the population is -3.85 ± 0.2×10^-8^ m^2^V^-1^s^-1^. ELS returns an EPM of -3.29 ± 2.7 ×10^-8^ m^2^V^-1^s^-1^ for these PS NPs. Measurement of these particles was replicated with single-particle and bulk measurements (Figure S8, Table S7 and S8). For both approaches, there is sample-to-sample variation in EPM. It is known that EPM is highly dependent on sample composition and not just the composition of the nanoparticle, so it is possible that sample-to-sample variation is caused by subtle changes in the suspension. Moving forward, this is also observed in the COOH-PS NPs.

The EPM of 198 nm and 196 nm COOH-PS were measured at the single-nanoparticle level and in bulk. 3D-SMART reports -4.08 ± 0.3 ×10^-8^m^2^V^-1^s^-1^and -3.91 ± 0.6 ×10^-8^m^2^V^-1^s^-1^, respectively (Fig. S9 and 10). Values reported from ELS for 198 nm and 196 nm COOH-PS are -4.08 ± 0.5 ×10^-8^ m^2^V^-1^s^-1^ and -1.13 ± 0.4 ×10^-8^ m^2^V^-1^s^-1^, respectively (Table S8 and S9). For the 198 nm COOH-PS NPs, the bulk and single-nanoparticle measurements return nearly identical values. Larger discrepancies arise for the 196 nm COOH-PS NPs. The magnitude of the EPM from ELS for 196 nm COOH-PS is consistently much lower than what is reported by 3D-SMART. We would assume that particles with only a 2 nm difference and the same surface functionalization would have very similar EPM as seen with 3D-SMART. Because of the lack of information about the distribution or correlated size of this population, it is difficult to discern the reason for this large deviation from what would be expected. Is it impossible to know if there is a subpopulation that is skewing the results in this way or if the EPM of each particle is nearly identical, leading to further questioning of the integrity of the sample. The independent measure of size on these samples does not give us much information, besides an accurate mean radius value that could suggest that the measured EPM is not due to aggregation (Table S13). The populations of 196 nm COOH-PS NPs highlight the importance of single-particle approaches to these types of problems.

Diffusion coefficient and electrophoretic mobility provide different approaches to characterizing populations of single nanoparticles. In Figure 4, it is apparent that the diffusion coefficient cannot statistically distinguish the difference in the populations of the 194 nm bare PS NPs and the 198 nm COOH PS NPs. However, when utilizing EPM of the populations and a Student’s t-test, it is determined that these populations are significantly different with a p value < .0001. This continues to hold true when pooling together replicates of each particle type. Figure S13 shows the distribution of diffusion coefficient and EPM for each particle for all replicates included in previous figures. Like the individual datasets provided in Figure 4, the combined data also show that this method can distinguish the difference between 194 nm PS NPs and 198 nm COOH-PS NPs using EPM. Including more data also allows us to distinguish 196 nm COOH-PS NPs from 198 nm PS NPs with EPM, unlike diffusion coefficient, with a p value < .0001. In the case of 194 nm PS NPs and 196 nm COOH-PS NPs, the diffusion coefficient is successful in determining the statistical difference between the population of (p < .05) but EPM can do so with greater certainty (p <.0001). Despite some inconsistencies across replicates, we maintain the ability to distinguish populations of nanoparticles only having a difference in (nominal) diameter of 2 nm using EPM, strengthening 3D-SMART’s characterization capabilities as a single particle method.

**Figure 4:**
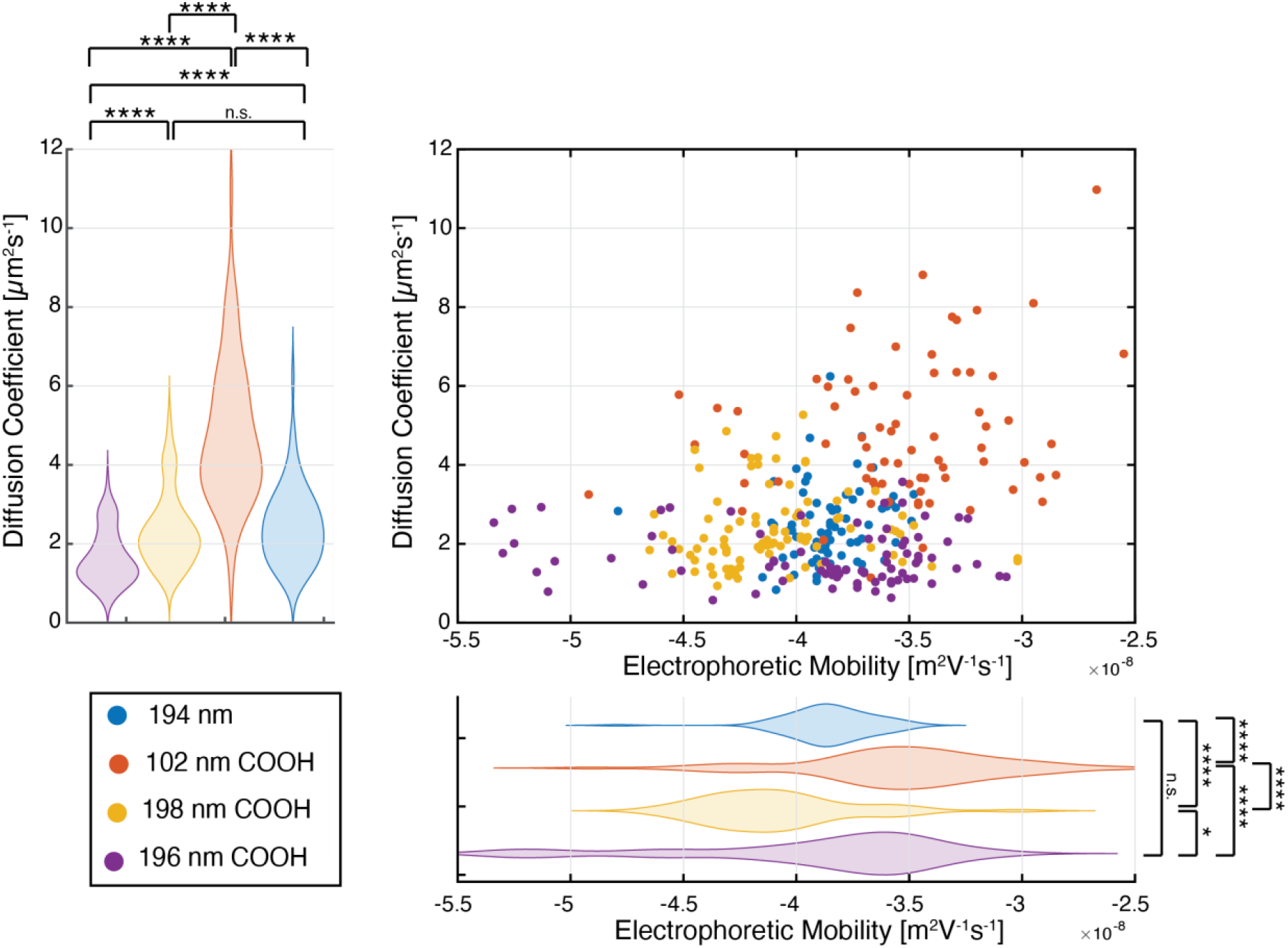
Electrophoretic mobility and diffusion coefficients of various sizes and surface functionalities of polystyrene nanoparticles. * = p < 0.05, **** = p < 0.0001, n.s. = not significant.

3D-SMART is also capable of tracking smaller particles to measure EPM. Another COOH-PS NPs used in this work is 102 nm COOH-PS NPs. The single-nanoparticle average EPM is -3.5 ± 0.4 ×10^-8^ m^2^V^-1^s^-1^ while ELS give us -4.43 ± 0.8 × 10^-8^ m^2^V^-1^s^-1^. Bulk data suggests that EPM magnitude would increase with decreasing size, while our single NP data suggests the opposite. The trend of EPM according to size is also not consistent throughout the findings of previously published single-nanoparticle EPM methods. Choi *et al*. saw EPM increase in magnitude with decreasing size while Oorlynck *et al*. saw the magnitude of EPM increase with size and ELS data substantiates their observed trends.

The findings reported in this current work can be validated using manufacturer specifications. The surface charge density is given for carboxyl-functionalized particles as a surface titer. For ∼100 nm and ∼200 nm COOH-PS NPs, the surface titers are 256 and 145, respectively. This gives a ratio of 1.8 for 100 nm to 200 nm COOH-PS NPs. Surface charge density is determined from 3D-SMART data as well. The effective charge of the double layer, Z_eff_, is calculated using Equation 6. Dividing Z_eff_ by the surface area of the spherical particle yields the surface charge density (Equation 11). For 102 nm COOH-PS NPs, the surface charge density is -1.1 ± 0.1 × 10^-3^ C/m^2^ and - 6.6 ± 0.3 × 10^-4^ C/m^2^ for 198 nm COOH-PS NPs. This gives a ratio of 1.7. The surface charge ratio of the experimental data closely matches the ratio calculated from the manufacturer’s specifications. Although we cannot establish a pattern for the size dependence of EPM for polystyrene nanoparticles overall, we are confident in the findings for the nanoparticles measured in this work.

Through single-nanoparticle tracking microscopy, 3D-SMART can report EPM of individual polystyrene nanoparticles and sample-to-sample variations seem to be a consequence of EPM being a highly sensitive parameter of spherical particle suspensions and not a decisive measure of the precision of 3D-SMART as bulk techniques are also vulnerable to these discrepancies.

### Ionic strength

There are many factors that affect the EPM of NPs in solution. The EPM of an individual NP reflects its environment, communicates the effective electric field in solution, availability of charge species in solution, and even solubility. One critical parameter that affects EPM is ionic strength. Electrolytes affect the local electric field strength and alter NP mobilities. For example, the addition of electrolytes such as NaCl into solution changes the ionic strength as well as the Debye length of the particles (Table S14). Here, ionic strengths ranging from 0.25-5 mM were used to test the effect of electrolytes on single-particle EPM. As shown in Figure 5A, an increase in ionic strength led to a consistent decrease in EPM for 196 nm COOH-PS NPs. This agrees with previous studies that show a decrease in EPM with increasing ionic strength.^23, 24^

**Figure 5:**
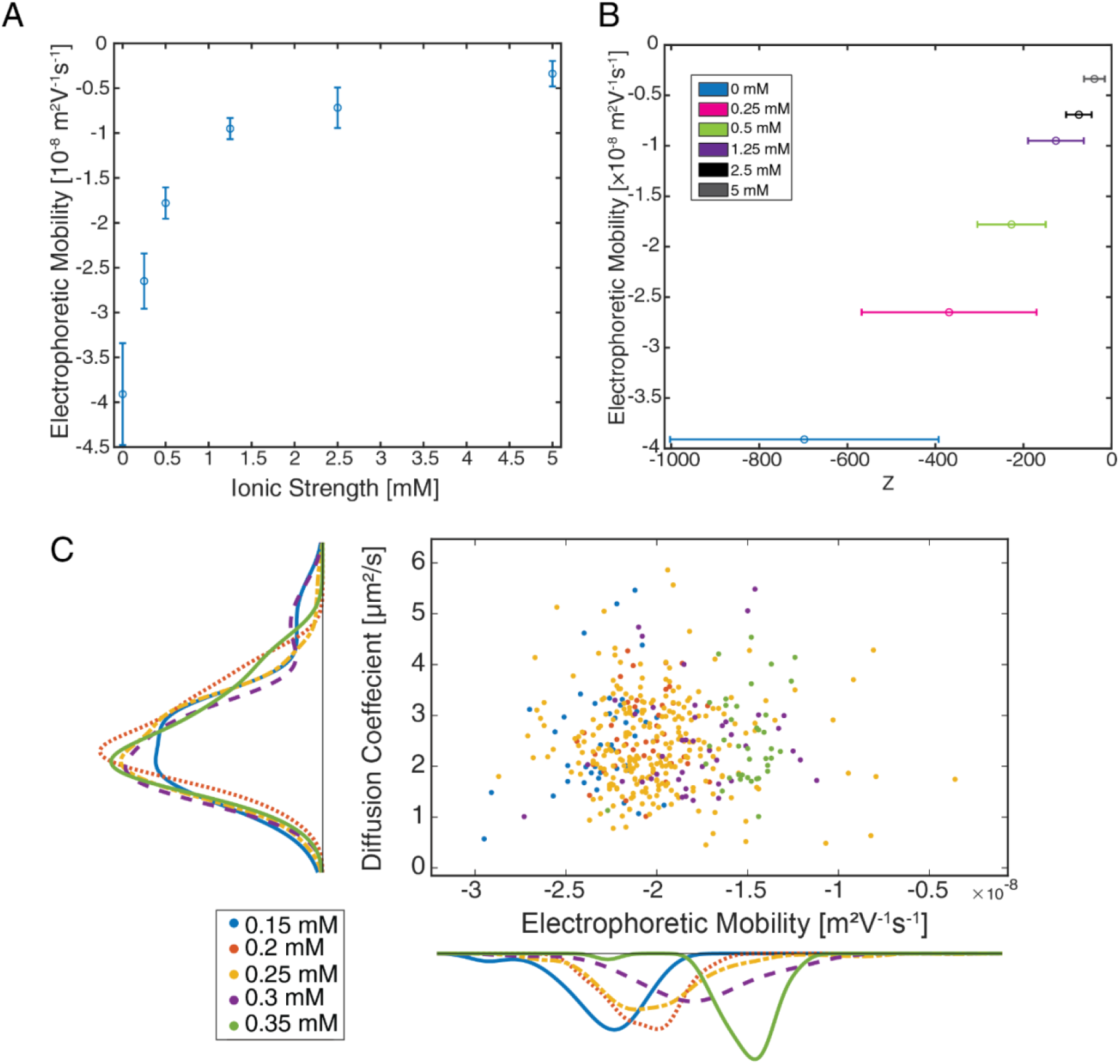
Ionic Strength dependence of single nanoparticle electrophoretic mobility. (A) Electrophoretic mobility of 196 nm carboxyl-functionalized polystyrene nanoparticles vs the ionic strength of 0-5 mM NaCl. (B) Electrophoretic mobility versus calculated charge number in NaCl solution for 196 carboxyl-functionalized polystyrene nanoparticles. Error bars represent the standard deviation of the population. (C) Electrophoretic mobility and diffusion coefficients of 194 nm unfunctionalized polystyrene nanoparticles with ionic strengths 0.15-0.35 mM at 0.05 mM increments.

From the Debye-Hückel model, it is known that the Debye length of a particle is dependent on the ionic strength of the solution (Equation 13). Particles in solution are surrounded by an uneven distribution of ions and counterions in solution that constitute the Debye-Hückel ion cloud. Along with the decrease in Debye length, we can also use the effective charge of the double layer (Z_eff_) as a function of EPM at different solvent conditions (Fig. 5B). The EPM is dependent on the Z_eff_ of the particle because it is closely tied to the surface charge of the nanoparticle. As the electrolyte concentration in solution increases (higher ionic strength), the counterions in solution become more tightly associated with the particle’s surface. The negative surface charge of the particle is increasingly screened by positive ions, causing the overall charge number to decrease as seen in Figure 5B. The valence charge number also reflects this change and can be calculated using Equation 14. The charge number is calculated for each individual NP based on the behavior seen in the trajectory as it is affected by the electric field. The lowered surface charge leads to a weaker reaction to the electric field, resulting in a lower EPM. We demonstrate here, that in addition to EPM, the charge number of individual NPs can be precisely measured by this new method. For the ionic strength range 0.25 mM – 5 mM, the mean charge numbers (± s.d.) were -369 ± 199, - 277 ± 77, -126 ± 63, -74 ± 29 and -39 ± 24, respectively as ionic strength increases. The mean uncertainty in charge number (± s.d.) for each population is 25 ± 9, 22 ± 6, 22 ± 7, 23 ± 10, and 24 ± 8. Across these ionic strengths explored, the valence charge number of individual nanoparticles can be determined within less than 30 units, demonstrating the precision of this method to determine the change of a single nanoparticle. 3D-SMART joins single molecule electrometry as a means of measuring the charge number of single particles in real time.^25^

Smaller shifts in the ionic strength were also explored to assess the precision of the method. NaCl was added to nanoparticle solutions in the ionic strength range 0.15 – 0.35 mM in intervals of 0.05 mM. In Figure 5C, EPM gradually shifted from left to right as the ionic strength increases. According to a student’s t-test the populations associated with 0.15, 0.25 and 0.35 mM are statistically different (p < 0.01). Using this method we can detect 0.1 mM shifts in ionic strength using EPM while maintaining accurate readings of particle size.

### Real-time particle manipulation with active-feedback field stimulation

A major difference between the active-feedback approach shown here and prior single-particle EPM measurements is the real-time measurement of the particle position which opens the possibility for real-time actuation of particles in solution. In previous work, the anti-Brownian electrokinetic (ABEL) trap has been able to trap single molecules in solution using electrokinetic feedback and microfluidics.^26-28^ The work presented here provides a foundation for single-particle trapping without the use of microfluidics and low field strengths, more closely mimicking the native environment for single molecules. To demonstrate this, a simple feedback scheme was applied to maintain the position of a single particle in solution in one dimension. An arbitrary position within the sample was selected. When the particle was measured to be on one side of the boundary, +2 V was applied to push the particle back to the selected position. On the other side of the boundary, -2 V was applied. The results of this simple confinement scheme are shown in Figure 6. Despite the simplicity of the employed feedback scheme, the 110 nm COOH-PS NP can be confined to within 2.95 μm of the center boundary as determined from the MSD of the trapped X-axis (Fig 6C and E), a dramatic confinement compared to the non-actuated Y-axis (Fig. 6D). The approach demonstrated here is a proof of principle that single-nanoparticles can be manipulated in solution in real-time in free solution and at low field strengths. With an appropriately designed sample chamber with field stimulation along all three axes, arbitrary control should be possible for a wide range of nanoscale objects in complex biological systems.

**Figure 6:**
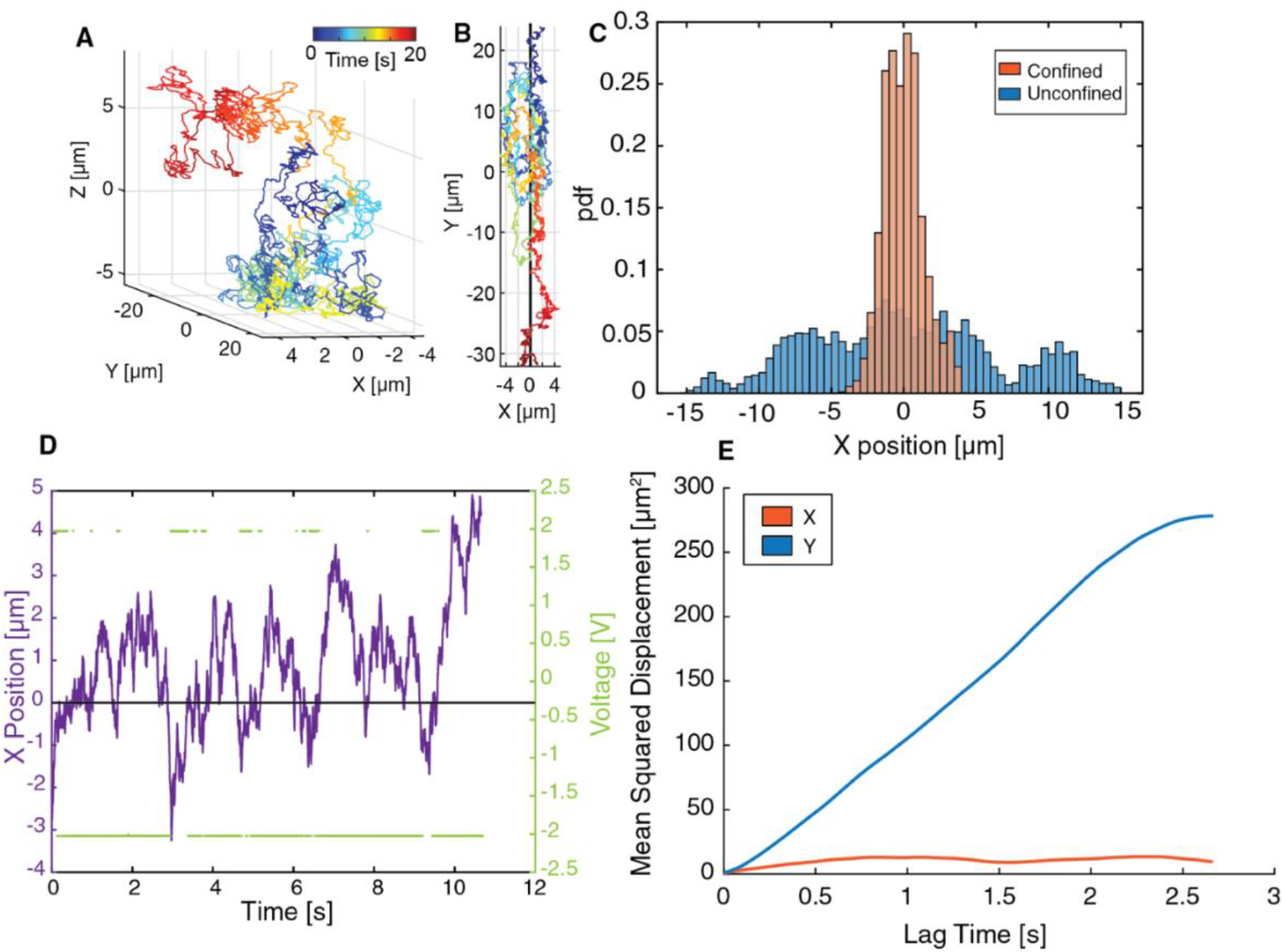
One-dimensional nanoparticle confinement via electrokinetic feedback. (A) 3D trajectory of 110 nm nanoparticle with electrokinetic feedback. (B) XY view of particle confinement trajectory using ± 2 V feedback. (C) X position distribution of unconfined and confined particles. (D) Particle confinement of X position (purple) and due to electrokinetic feedback (green) with a 40 µm boundary. (E) 1D MSD of directed motion along X-axis (orange) and free diffusion along Y axis (blue).

## Conclusion

The single-nanoparticle EPM demonstrated here has a myriad of future potential applications, most of which involve measuring EPM in complex environments. Future expansion of this new technique to biological environments will require overcoming several potential obstacles. The first is the conductivity of buffer solutions used or present in most biological systems (e.g. phosphate buffered saline). As the buffer conductivity increases, higher currents will be required to reach the same field strength. Careful consideration of how the present electrolytes contribute to redox reactions and heating of solution at increased ionic strengths will be critical. The seconds issue going forward is the decreased observation time of 3D-SMART as the diffusive speed of the probe increases. In its current state 3D-SMART can track ∼1 kB double-stranded DNA for several seconds.^29^ This does not meet the 20-second limit corresponding to a threshold demonstrated above. As previously mentioned, a higher field strength can eliminate the need for a time cut off but it will decrease the observation time even further. Further exploration of the optimal field strength and frequency will be beneficial in the future as species of interest become smaller and diffuse more rapidly.

In the future, it should also be possible to take advantage of 3D-SMART’s ability to be paired with concurrent imaging techniques (3D-TRIM)^18^ and measure how the EPM of nanoparticles, the protein corona, or even viruses change as they come in close proximity to cells. It should also be possible to measure the surface charge density of nanoparticles in these environments (Table S16). Surface charge density has been used in previous studies to assess nanoparticle as viable drug and gene carriers.^30-32^ Real-time measurement of surface charge density and nanoparticle behavior in a cellular environment cold give valuable insight into drug delivery and therapeutic design. Without reliance on microfluidics, translating this method to accommodate live tissue can be done with few adjustments to the experimental set up. However, this would require meticulous optimization as tissue could distort the electric field.

Here, we have demonstrated that 3D-SMART enables in situ electrophoretic mobility determination of single nanoparticles, with single nanoparticle charge calculations to within 30 charge units. Importantly, the size of the nanoparticle is simultaneously extracted. This means that any level of aggregation within solution, which in ensemble methods skews the reported values for particles in solution, is not a problem for this method as aggregates are easily discerned via MSD analysis. In contrast to prior work on single-nanoparticles EPM extraction, the work described above reports a more accurate particle diameter, can use long observation times to monitor changes and does not require microfluidics, making the expansion to complex biological samples possible (Table S1). Furthermore, the real-time nature of 3D-SMART makes active electrophoretic confinement of a single nanoparticle possible, shown here for the first time without the use of microfluidics. This suggests that 3D confinement and manipulation will be possible in future work with an appropriately constructed sample chamber. Being able to precisely control and manipulate nanoscale objects will open new lines of investigation across the biological sciences.

## Methods

### Application of Electric Field

Using a closed bath imaging chamber with field stimulation (Warner Instruments RC-21BRFS), an electric potential was applied across the 6.3 mm gap between two platinum wires (Figure 1B). The applied electric field strength in this work was between -3.2 V/cm and +3.2 V/cm. Voltage was supplied by the FPGA utilized for particle tracking and control via National Instruments LabVIEW software. The direction of the field corresponds to the X axis of the 3D tracking volume.

### 3D-SMART

As previously published, 3D-SMART uses a “lock on” algorithm to track freely diffusing nanoparticles in real time. A focused laser spot (488 nm) scans a 5×5 tracking volume in a knight’s tour pattern using two electro-optical deflectors (EODs) in XY, while a tunable acoustic gradient (TAG) lens scans Z in a sine wave pattern (Fig 1C). The EODs deflect the KT spending 20 μs per spot while the TAG has a 14 μs period for the generated sine wave. Photon arrival times are collected by an avalanche photodiode (APD) and used to estimate particle position in XYZ through a Kalman filter-based algorithm.^17^ A piezoelectric stage and 2D galvo scanning mirror compensate for particle movement within the tracking volume (1 μm × 1 μm × 2 μm), ensuring that the particle remains within the field of view. Real-time particle coordinates correspond to the stage and galvo positions allowing generation of 3D trajectories (Fig 1D). The EODs, TAG, APD, stage and galvo are controlled via FPGA. Post-processing of tracking data is performed in MATLAB. All nanoparticles were diluted to ∼1 pM to avoid the possibility of multiple particles in the observation volume.

### Mean Squared Displacement

Three-dimensional mean squared displacement (MSD) of individual nanoparticle trajectories were calculated using 3D position information and equation S1:

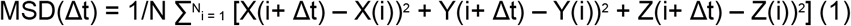

N is the number of data points for the calculation and Δt is the lag time. The 3D coordinates of the particle are represented by X, Y, and Z. Linear fitting of the MSD vs lag time yield the diffusion coefficient of the nanoparticle due to the Stokes-Einstein equation:

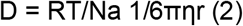

where R is the gas constant, Na is Avogadro’s number, η is the dynamic viscosity of the solution and r is the radius of the NP. The NP exhibits three-dimensional Brownian motion in the absence of any external forces in solution so the behavior of MSD vs lag time plot is expected to be linear.

One-dimensional position data was used to calculate 1D MSDs in a similar manner, using a modification of equation 1 for a single dimension.

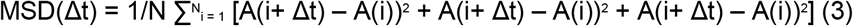

A represents the particle’s one-dimensional position (X, Y, or Z). The one-dimensional coordinate replaces the other two dimensions to calculate MSD. Linear fitting of MSD vs. lag time remains the same. Diffusion coefficients are extracted from the MSD vs. lag time plots for the non-actuated axes. The X coordinates of field-stimulated NPs do not return accurate diffusion coefficients or hydrodynamic radii, as they are not undergoing Brownian motion.

### Size Determination

Radii of NPs under field stimulation only factor in the diffusion coefficients of the non-actuated axes (Y and Z). Two-dimensional MSDs are used for size characterization.

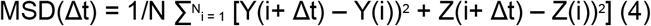

The diffusion coefficient is determined by linear fitting of the MSD vs. lag time plot, and the Stokes-Einstein equation yields the hydrodynamic radius.

### Electrophoretic mobility

The electrophoretic mobility can be related to the experimentally measured drift velocity (u) and the applied electric field (E) via Equation 5. Electrophoretic mobility can also be expressed in terms of voltage and chamber length (d) via Equation 6.

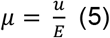

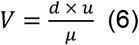

The displacement of the particle along the X-axis of the tracking volume is calculated based on the coordinate data recorded via 3D-SMART to calculate the electrophoretic mobility of the particle. The voltage applied by the FPGA and the distance between the platinum electrodes (d) were used to calculate *E*. The probability density of observing a particle displacement, Δ*x*, after a time τ with diffusion coefficient *D* and drift velocity *u*, follows a Gaussian probability distribution.

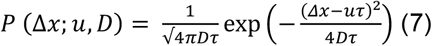

Under the conditions where a constant drift velocity is expected (under constant voltage), the mean displacement for a given time τ is sufficient to extract the EPM. However, this will yield a EPM of zero for an oscillating potential. To account for this, we apply a maximum likelihood approach to extract the EPM give a time-varying potential. The log-likelihood, *L*, of a particular *u*_*i*_ and D given a set of observed displacements Δ*x*_*i*_ is

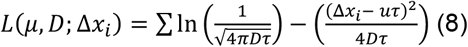

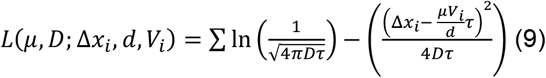

Finding the maximum of Equation 9 can yield the maximum likelihood estimation (MLE) of both µ and *D*. In the work described below, the value of *D* is calculated from the mean-squared displacement along the non-actuated axis (y) and a one-dimensional log-likelihood is used to find the value of the drift velocity. The MLE and 95% confidence interval are reported for all data.

### Zeta Potential

The zeta potential (ξ) can be calculated based on the drift velocity (*u*), solvent viscosity (η), chamber width (d), applied voltage and relative and vacuum permitivities (ε_r_ and ε_0_).

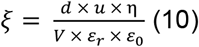

### Effective Surface Charge

The effect charge of the double layer (Z_eff_) is given by the mobility (μ_E_), viscosity and Stoke’s radius (R_s_).

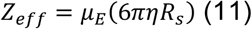

### Surface Charge Calculation

The combination of experimentally determined surface charge (Z_eff_, Eq. 8) and manufacturer-reported surface area (*A*) of the NP makes the determination of the surface charge density possible for spherical nanoparticles.

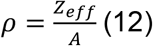

### Debye Length

The Debye length (*κ*^−1^) is given by:

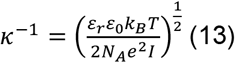

Here *N*_*A*_ is Avogadro’s number, *e* is the elementary charge, and *I* is the ionic strength. Higher ionic strength thins the electric double layer.^23^

### Valence Charge number

The valence charge number is calculated by dividing the effective charge of the double layer (Z_eff_) by the elementary charge of an electron (e).

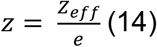

### DLS and ELS measurements

Samples prepared for DLS were measured using a Malvern ZetaSizer Ultra Red. Technical replicates were diluted 1000x from stock solution of each nanoparticle, rinsing the cuvette with deionized water in between measurements.

## Acknowledgments

We acknowledge financial support from the National Institute of General Medical Sciences of the National Institutes of Health under award number R35GM124868 (K.D.W.). This material is based upon work supported by the National Science Foundation Graduate Research Fellowship Program under Grant No. DGE 2139754 (A.J.). This work was performed in part at the Duke University Shared Materials Instrumentation Facility (SMIF) (RRID:SCR_027480), which is supported by the National Science Foundation (award number ECCS-2025064).

