## Supporting Information for "Revision: Single-nanoparticle electrophoretic mobility determination and trapping using active feedback 3D tracking"

**Alexis Johnson and Kevin D. Welsher**
Department of Chemistry, Duke University, Durham, NC 27708, USA

### Materials

*Nanoparticles*

Green fluorescent carboxyl-functionalized polystyrene (102 nm/110 nm/196 nm/198 nm) and unfunctionalized polystyrene (194 nm) nanoparticles were purchased from Bangs Laboratories, Inc.

*Sample chamber*

Slotted Bath with field stimulation (RC-21BRFS) purchased from Warner Instruments.

**Supplemental Figures**

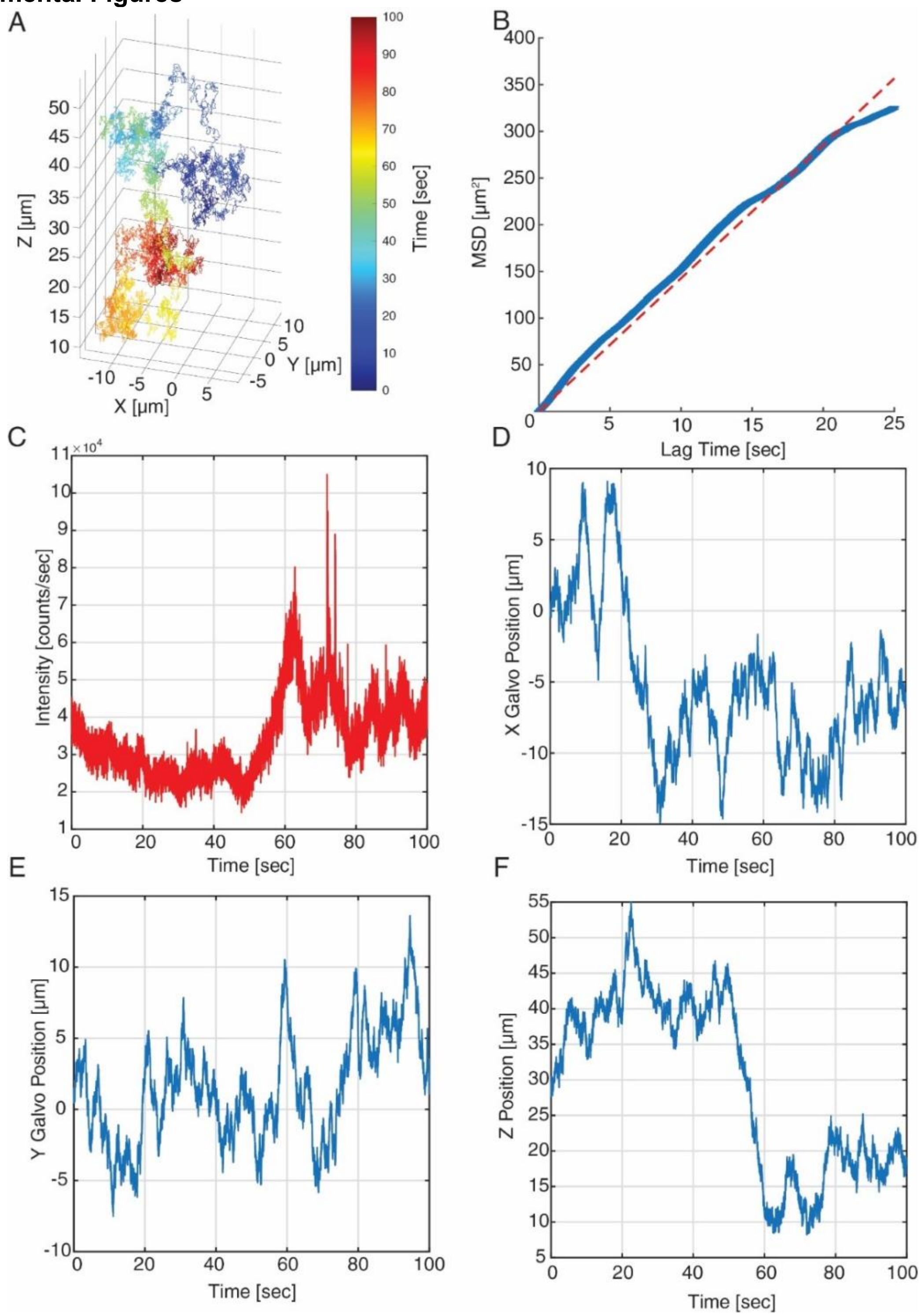

**Figure S1:** 3D-SMART tracking of individual nanoparticles. (A) 3D Trajectory. (B) 3D MSD. (C) Detected intensity. (D) X position readout from Galvo scanning mirror (E) Y position from Galvo scanning mirror. (F) Z stage position.

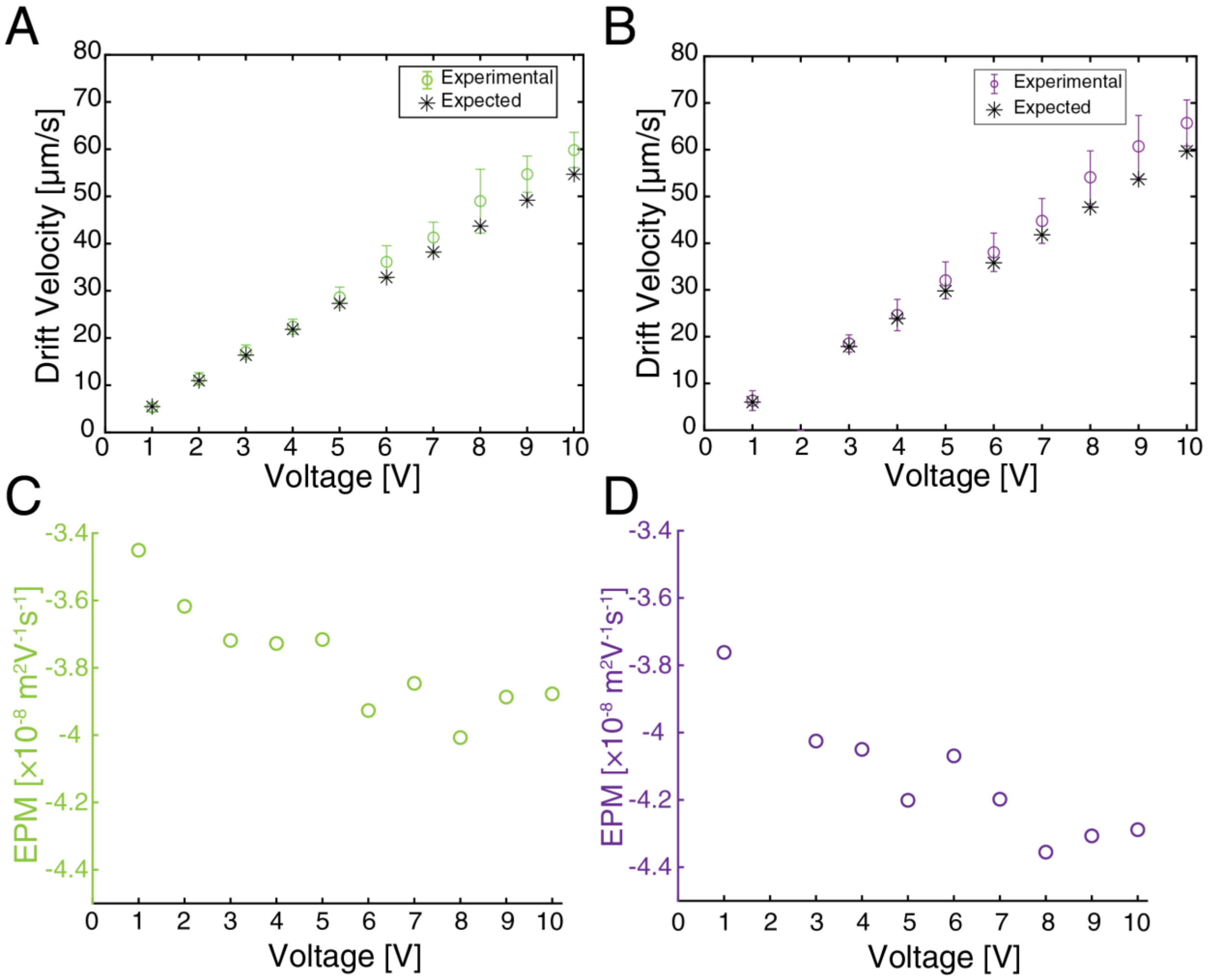

**Figure S2:** Experimental and expected drift velocity of 198 and 102 nm carboxyl functionalized polystyrene nanoparticle in water. (A) Expected and experimental drift velocity of 198 nm carboxyl functionalized nanoparticles under the full voltage range of the FPGA (1 –10 V). (B) Expected and experimental drift velocity of 102 nm carboxyl functionalized nanoparticles under the full voltage range of the FPGA. (C) Measured EPM of 198 nm carboxyl functionalized NPs at 1-10 V (D) Measured EPM of 102 nm carboxyl functionalized NPs at 1-10 V.

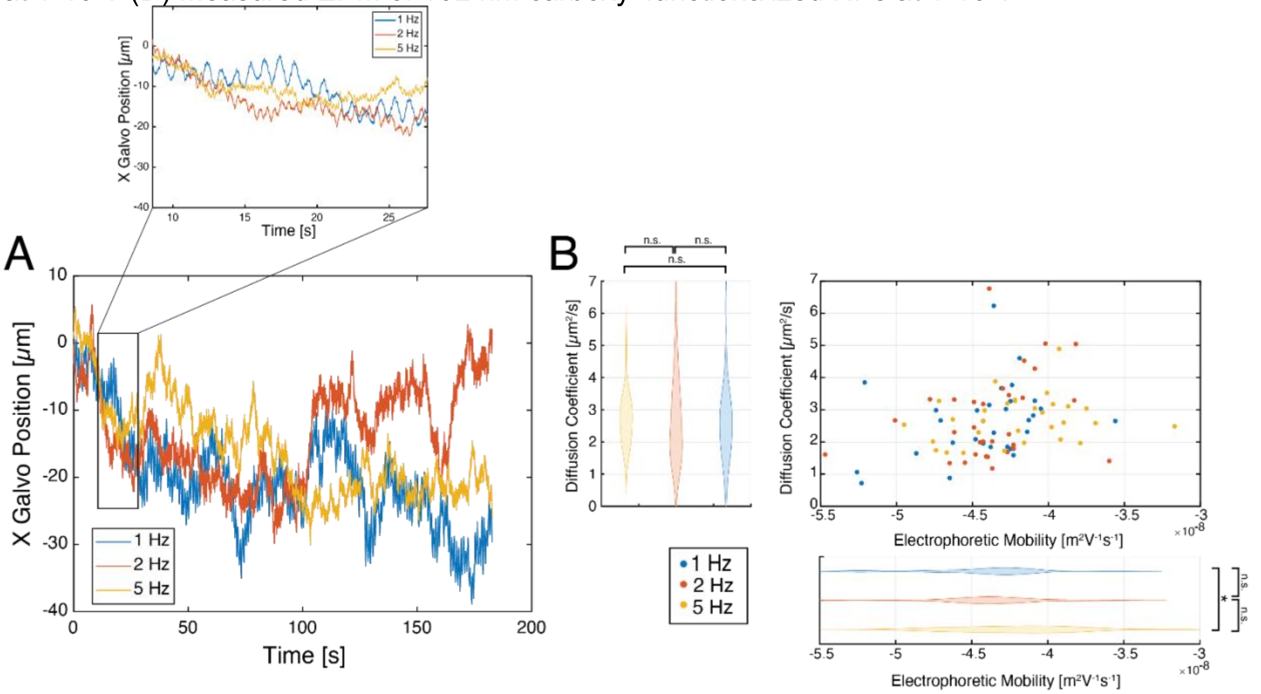

**Figure S3:** Frequency variation of 2 V oscillating electric field. (A) X galvo position over time at frequencies 1, 2 and 5 Hz of 198 nm COOH PS NPs at 2 V. Inlet: Zoomed in region of A. (B) Diffusion coefficient and electrophoretic mobility of 198 nm COOH PS NPs in water at 2 V and frequencies 1, 2 and 5 Hz. n.s. = not significant, * = $p < 0.05$.

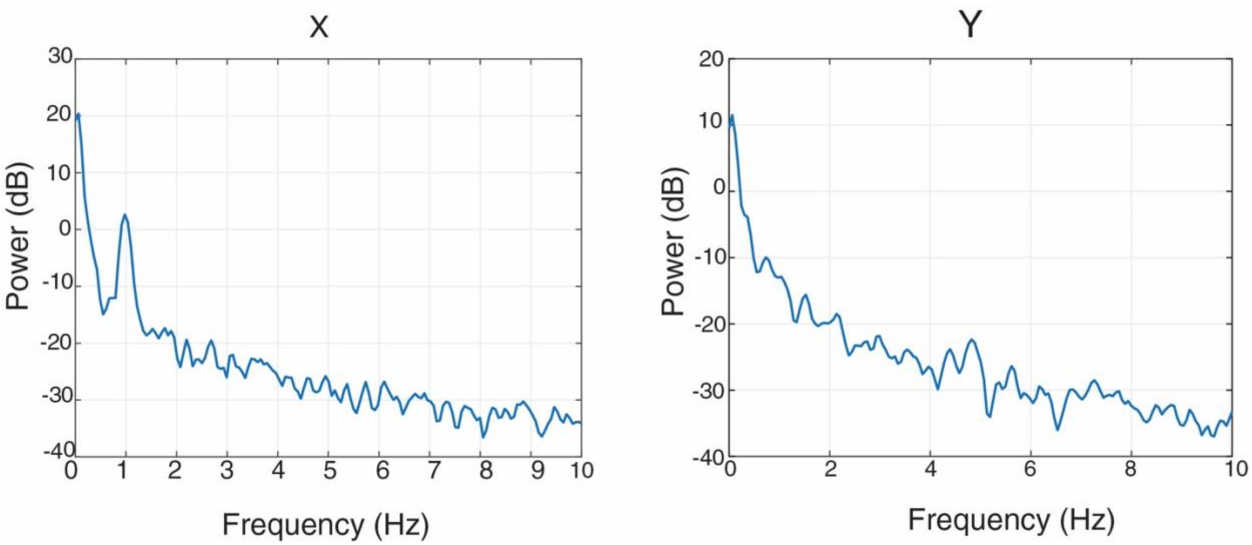

**Figure S4**: Welch's power spectral estimate analysis of actuated (X) and non-actuated (Y) axes of trajectory under oscillating electric field.

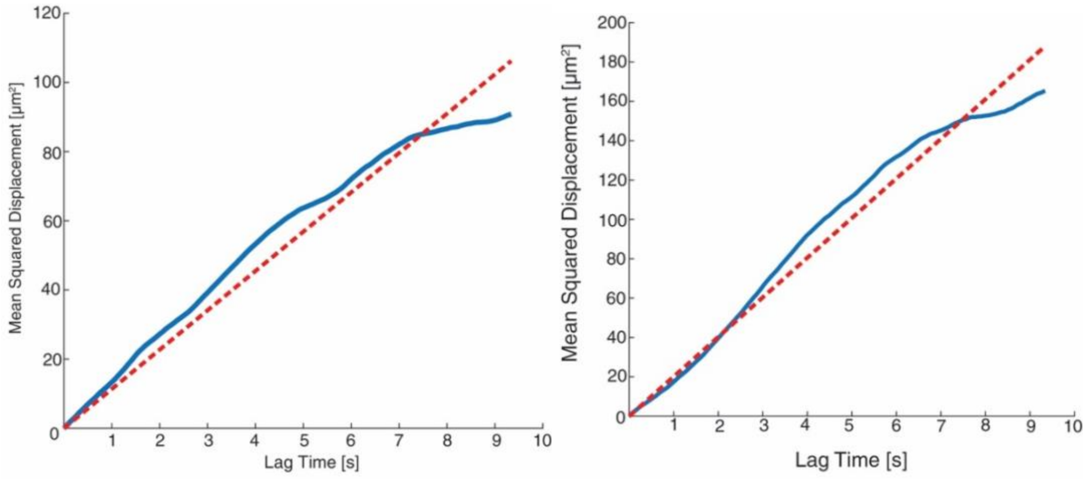

**Figure S5**: Mean squared displacement for Y (left) and Z (right) axes from Figure 2 oscillating electric field trajectory.

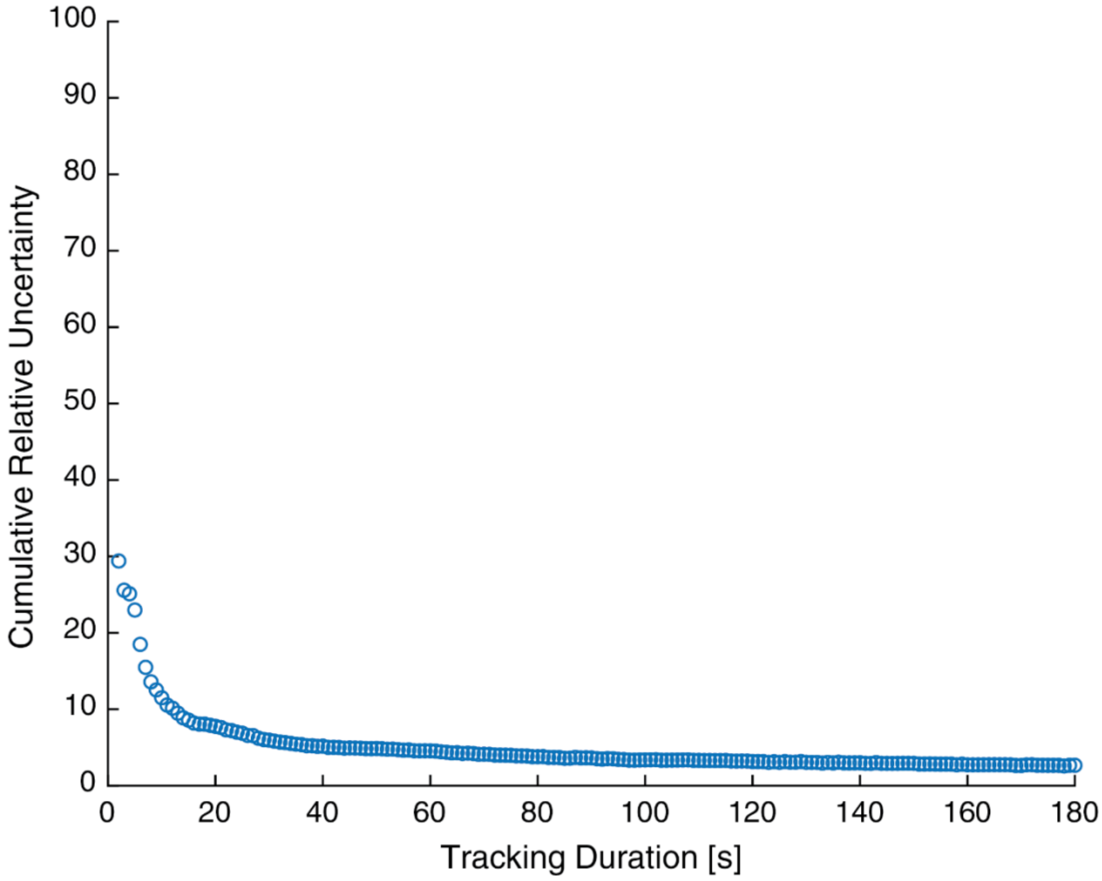

**Figure S6:** Cumulative Relative Uncertainty of EPM as a function of time for a single 102 nm carboxyl functionalized polystyrene nanoparticle.

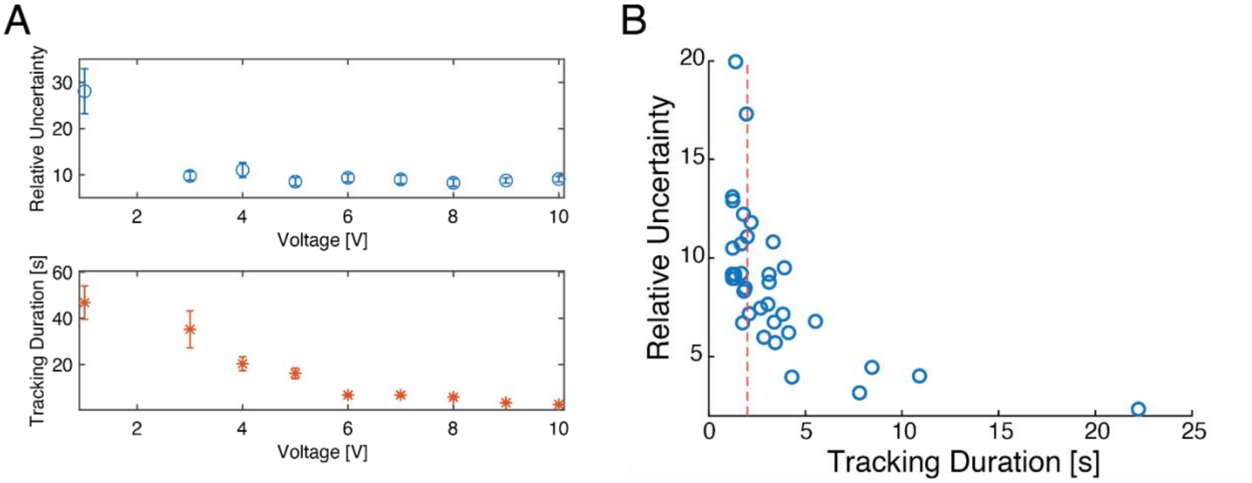

**Figure S7:** EPM precision vs voltage and tracking duration. (A) Top: Mean relative uncertainty (95% confidence interval). Error bars denoting standard error of the mean. N = 30. Bottom: Mean tracking duration of 102 nm COOH PS NPs in voltage range 1-10 V. Error bars denoting standard error of the mean. N = 30. (B) Relative uncertainty vs tracking duration of 102 nm COOH PS NPs at 10 V. Dashed red line denoting 2 seconds of tracking.

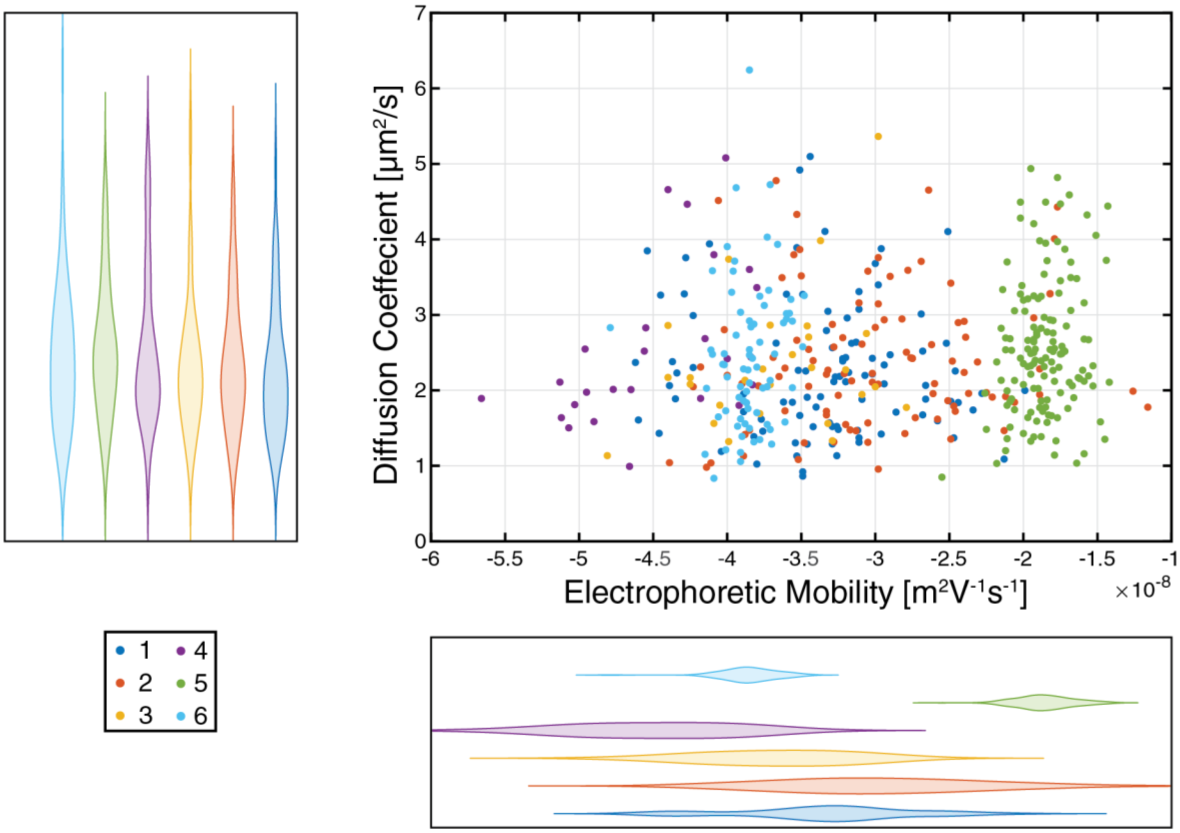

**Figure S8:** Scatter plot of single nanoparticle diffusion coefficient and electrophoretic mobility of 194 nm polystyrene nanoparticles in water. N = 97, 98, 27, 136, 73, respectively.

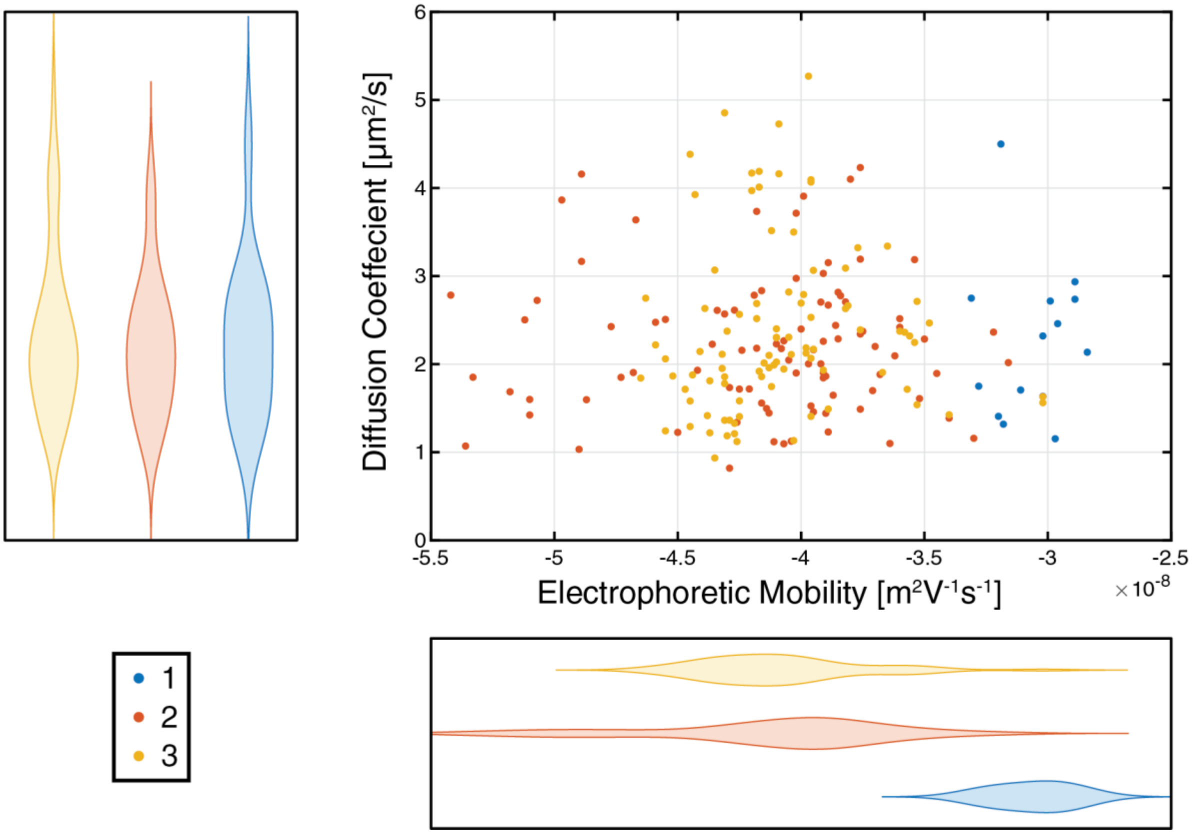

**Figure S9:** Scatter plot of single nano particle diffusion coefficient and electrophoretic mobility of 198 carboxyl functionalized nm polystyrene nanoparticles in water. N = 14, 92, 92, respectively.

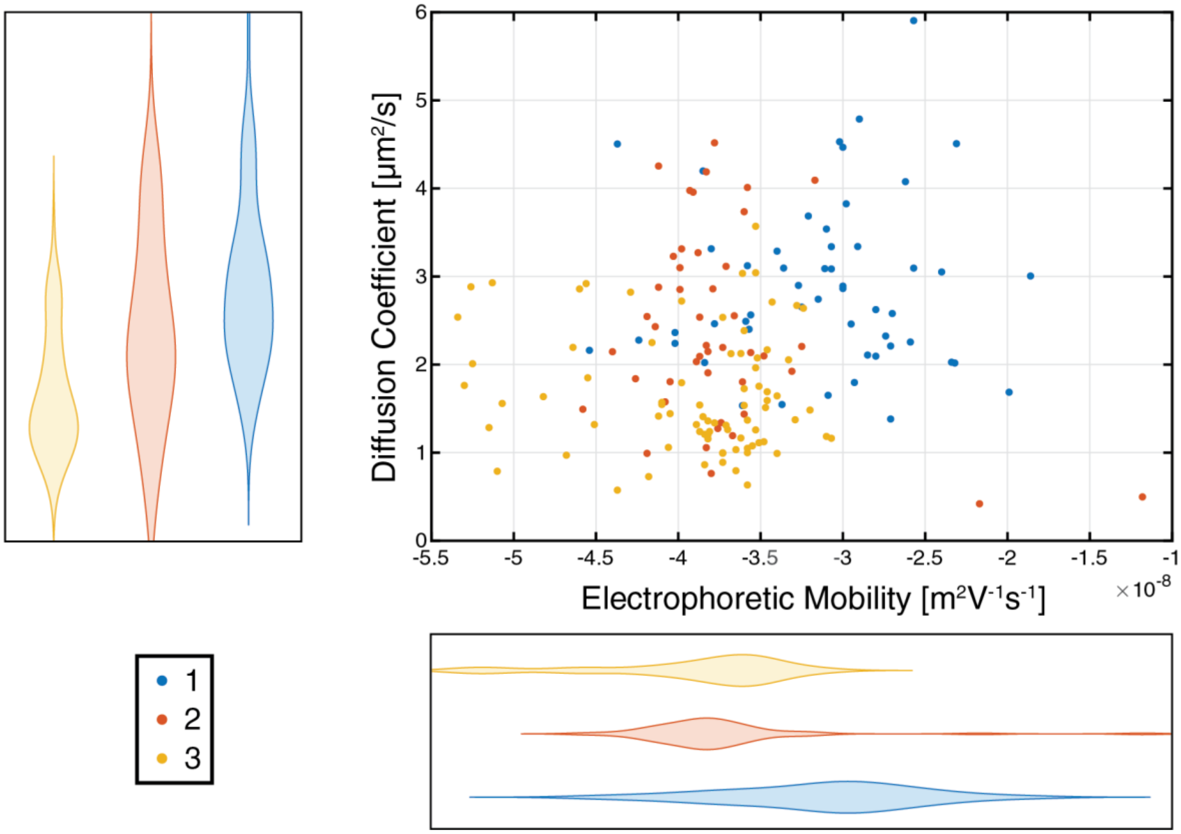

**Figure S10:** Scatter plot of single nanoparticle diffusion coefficient and electrophoretic mobility of 196 nm carboxyl functionalized polystyrene nanoparticles in water. N = 52, 45, 79, respectively.

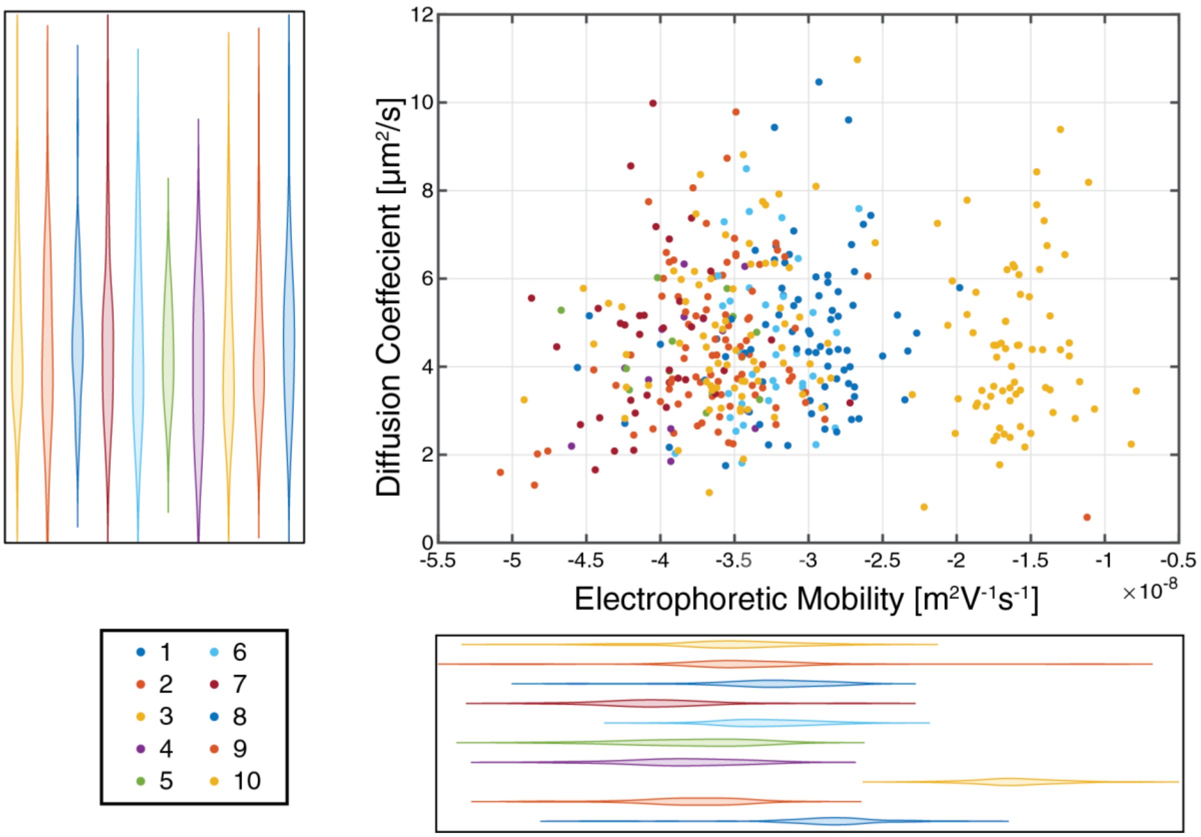

**Figure S11:** Scatter plot of single nanoparticle diffusion coefficient and electrophoretic mobility of 102 nm carboxyl functionalized polystyrene nanoparticles in water. N = 46, 43, 68, 13, 14, 34, 35, 30, 45, 73.

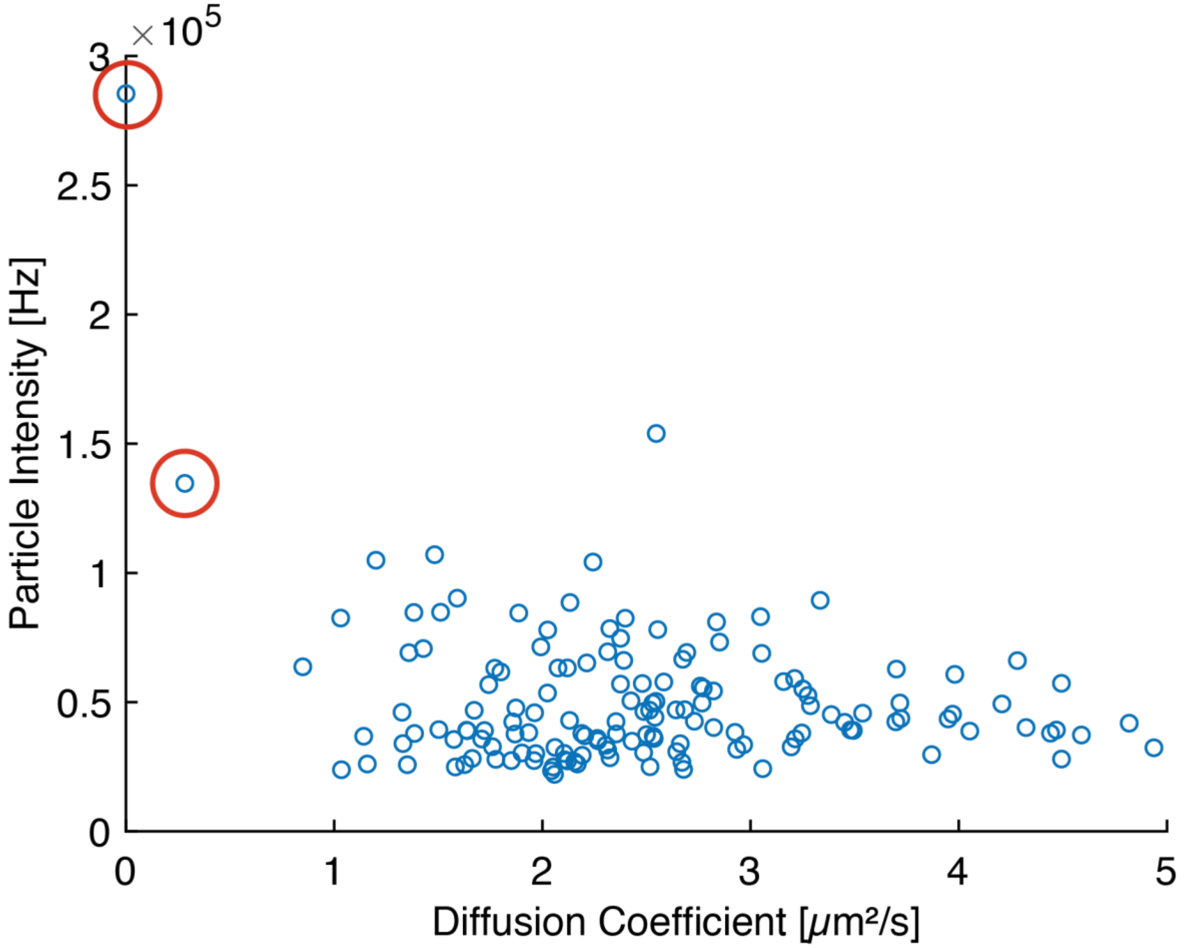

**Figure S12:** Determination of aggregate by particle intensity and diffusion coefficient. Red circle denotes aggregates.

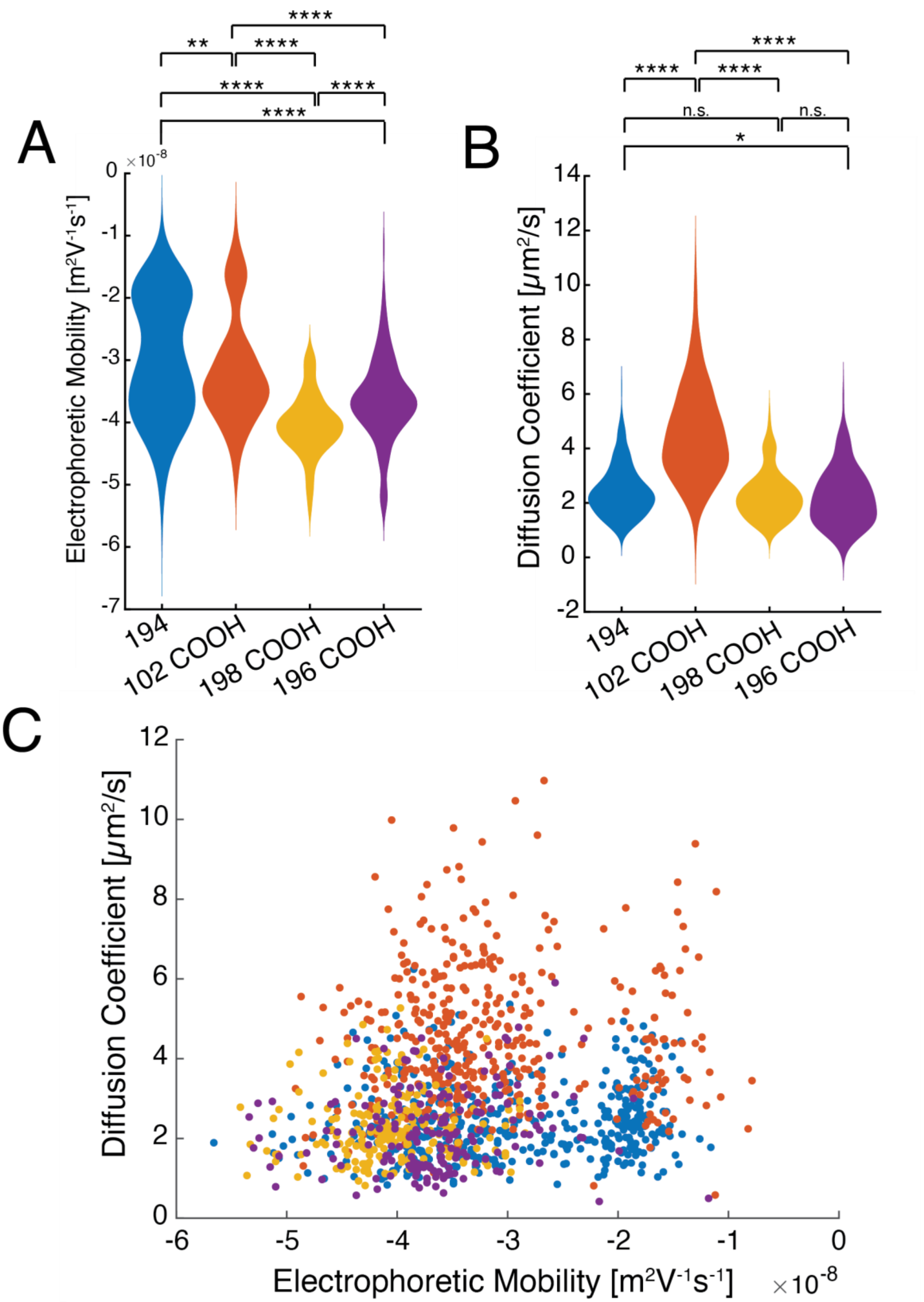

**Figure S13:** Mean Electrophoretic mobility and diffusion coefficient. (A) Distribution of electrophoretic mobility for all nanoparticle types. (B) Distribution of diffusion coefficient for all nanoparticle types. (C) Scatter plot of corresponding diffusion coefficient and electrophoretic mobility. Colors correspond to A and B.

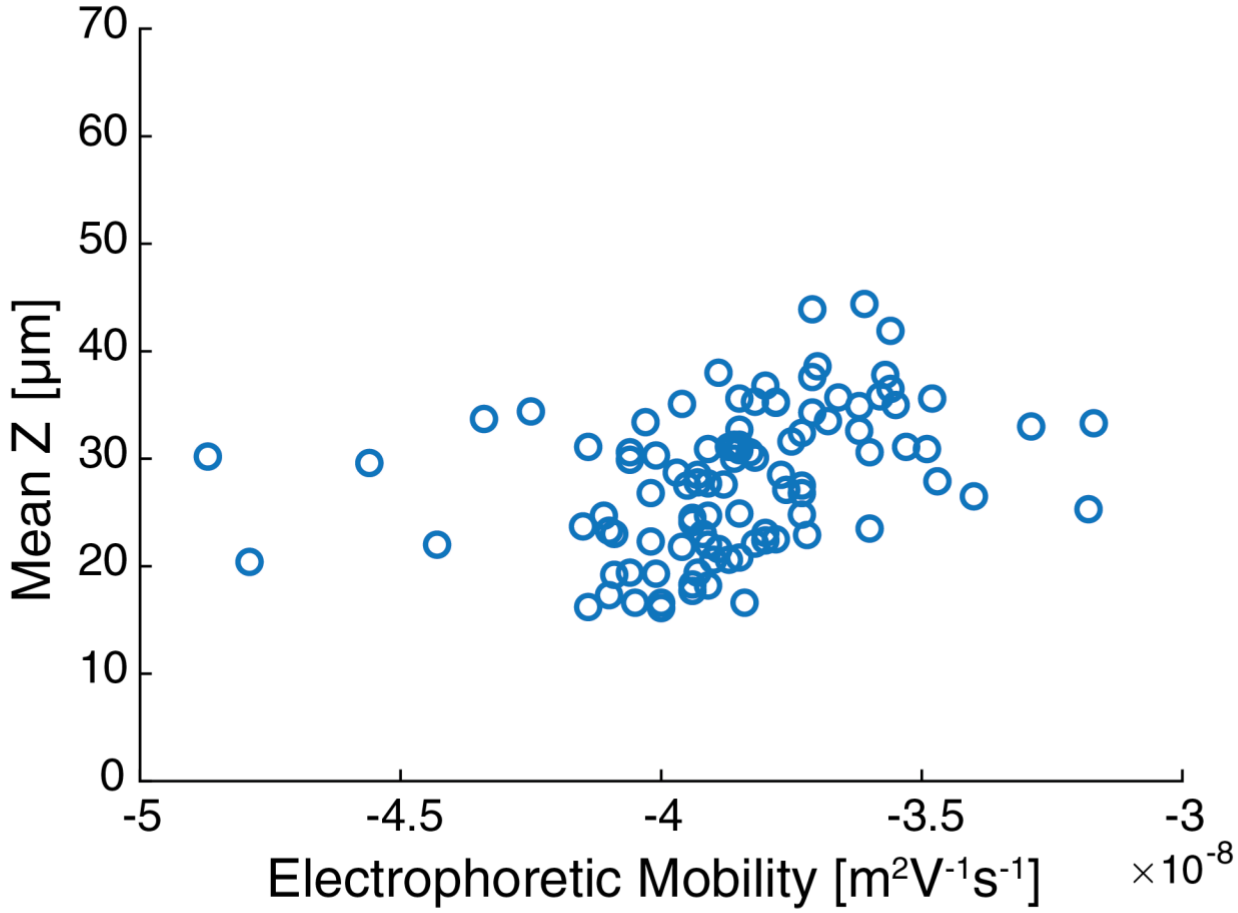

**Figure S14:** Mean Z position of 194 nm bare PS NPs in water vs electrophoretic mobility.

**Supplementary Tables**

Table S1: Single particle mobility determination method comparison

| Parameter | 3D-SMART | Choi *et. Al.* (2024) | Oorlynck *et. al.* (2023) |
|---|---|---|---|
| **Chamber dimensions** | 18 x 6.3 x 2.3 mm | 300 µm x 11 mm x 3 mm | 10 mm 6 mm x 100 µm |
| **Electrode spacing** | 6.3 mm | 100 µm | 6 mm |
| **Electrode type** | Platinum | Gold plated | Cr/Au coated glass slide |
| **Applied Voltage** | 1-10 V | 0.75, **1**, 1.5 V | 80 V |
| **Applied Electric field** | 3.2 V/cm | 75, **100**, 150 V/cm | 133 V/cm (167 is the max) |
| **Wave type** | Sine | **Sine** and square | Sine |
| **Frequency** | 1 Hz | **50**, 75 Hz | 300 Hz |
| **Z range** | 70 µm | 8 µm | 1.4 µm |
| **Observation time** | Up to 3 minutes | 1 second | 0.75 seconds |
| **Particle diameter** | 100, 200 nm | 530 nm, 840 nm, 1 µm, 2 µm | 100 nm, 500 nm |
| **Temporal resolution** | 210 µs | 1000 fps (1 ms) | 3300 fps (300 µs) |
| **Highest reported precision (95% confidence interval*)** | ~9% (100 nm)<br>~5% (200 nm) | ~4% (500 nm) | ~15% (100 nm)<br>~9% (500 nm) |
| **Solvent** | Water | 0.1 mM PBS buffer | Water |

**Table S2:** Mean Electrophoretic mobility and diffusion coefficient

| **Particle** | **3D-SMART** | | **ELS** | |
|---|---|---|---|---|
| | EPM (× $10^{-8}$ $m^2V^{-1}s^{-1}$) | Diffusion Coefficient ($\mu m^2 s^{-1}$) | EPM (× $10^{-8}$ $m^2V^{-1}s^{-1}$) | Diffusion Coefficient ($\mu m^2 s^{-1}$) |
| **194 BARE** | -2.98 ± 1 | 2.43 ± 0.91 | -3.72±0.20 | 2.50±0.04 |
| **198 COOH** | -4.04 ± 0.5 | 2.29 ± 0.87 | -3.79±0.24 | 2.56±0.02 |
| **196 COOH** | -3.64 ± 0.7 | 2.21 ± 1.0 | -1.09±0.03 | 2.22±0.20 |
| **102 COOH** | -3.20 ± 0.9 | 4.57 ± 1.7 | -4.26±0.38 | 4.58±0.14 |

Table S3: Mean electrophoretic mobility and diffusion coefficient of 194 nm unfunctionalized nanoparticles by 3D-SMART.

| **194 Replicate** | EPM ($10^{-8}$ $m^2V^{-1}s^{-1}$) | Diffusion Coefficient ($\mu m^2/s$) | Diameter (nm) |
|---|---|---|---|
| **1** | -3.34 ± 0.59 | 2.26 ± 0.88 | 219 ± 83 |
| **2** | -2.98 ± 0.69 | 2.39 ± 0.88 | 205 ± 79 |
| **3** | -3.67 ± 0.50 | 2.34 ± 0.88 | 205 ± 69 |
| **4** | -4.49 ± 0.51 | 2.54 ± 1.03 | 194 ± 74 |

| | | | |
|---|---|---|---|
| **5** | -1.86 ± 0.19 | 2.56 ± 0.88 | 190 ± 75 |
| **6** | -3.85 ± 0.19 | 2.45 ± 0.96 | 202 ± 82 |

Table S4: Mean electrophoretic mobility and diffusion coefficient of 198 nm carboxyl-functionalized nanoparticles by 3D-SMART.

| **198 Replicate** | EPM ($10^{-8}$ $m^2V^{-1}s^{-1}$) | Diffusion Coefficient ($\mu m^2/s$) | Diameter (nm) |
|---|---|---|---|
| **1** | -3.06 ± 0.14 | 2.25 ± 0.85 | 216 ± 76 |
| **2** | -4.14 ± 0.50 | 2.21 ± 0.77 | 219 ± 82 |
| **3** | -4.08 ± 0.32 | 2.37 ± 0.95 | 208 ± 76 |

Table S5 Mean electrophoretic mobility and diffusion coefficient of 196 nm carboxyl-functionalized nanoparticles by 3D-SMART.

| **196 Replicate** | EPM ($10^{-8}$ $m^2V^{-1}s^{-1}$) | Diffusion Coefficient ($\mu m^2/s$) | Diameter (nm) |
|---|---|---|---|
| **1** | -3.12 ± 0.58 | 2.92 ± 0.95 | 167 ± 53 |
| **2** | -3.75 ± 0.54 | 2.46 ± 1.05 | 238 ± 180 |
| **3** | -3.91 ± 0.57 | 2.79 ± 1.02 | 202 ± 83 |

Table S6: Mean electrophoretic mobility and diffusion coefficient of 102 nm carboxyl-functionalized nanoparticles by 3D-SMART.

| **102 Replicates** | EPM ($10^{-8}$ $m^2V^{-1}s^{-1}$) | Diffusion Coefficient ($\mu m^2/s$) | Diameter (nm) |
|---|---|---|---|
| **1** | -2.90 ± 0.45 | 4.80 ± 1.77 | 102 ± 41 |
| **2** | -3.80 ± 0.35 | 4.47 ± 1.65 | 109 ± 39 |
| **3** | -1.60 ± 0.29 | 4.39 ± 1.75 | 117 ± 64 |
| **4** | -3.84 ± 0.34 | 4.03 ± 1.45 | 123 ± 50 |
| **5** | -3.81 ± 0.39 | 4.37 ± 0.95 | 103 ± 22 |
| **6** | -3.28 ± 0.27 | 4.55 ± 1.75 | 110 ± 45 |
| **7** | -4.03 ± 0.39 | 4.60 ± 1.80 | 108 ± 34 |
| **8** | -3.26 ± 0.38 | 4.71 ± 1.51 | 101 ± 34 |
| **9** | -3.46 ± 0.55 | 4.40 ± 1.75 | 126 ± 106 |
| **10** | -3.53 ± 0.43 | 4.83 ± 1.77 | 102 ± 48 |

Table S7: ELS determination of electrophoretic mobility of 194 nm unfunctionalized nanoparticles.

| **194 Replicate** | EPM ($10^{-8}$ $m^2V^{-1}s^{-1}$) |
|---|---|
| **1** | -3.81 ± 0.55 |
| **2** | -3.86 ± 0.65 |

| | |
|---|---|
| **3** | -3.79 ± 0.63 |
| **4** | -3.78 ± 0.54 |
| **5** | -3.82 ± 0.63 |
| **6** | -3.29 ± 2.65 |

Table S8: ELS determination of electrophoretic mobility of 198 nm carboxyl functionalized nanoparticles.

| **198 COOH Replicate** | EPM ($10^{-8}$ $m^2V^{-1}s^{-1}$) |
|---|---|
| **1** | -3.57 ± 0.79 |
| **2** | -4.09 ± 0.45 |
| **3** | -4.08 ± 0.52 |

Table S9: ELS determination of electrophoretic mobility of 196 nm carboxyl functionalized nanoparticles.

| **196 COOH Replicate** | EPM ($10^{-8}$ $m^2V^{-1}s^{-1}$) |
|---|---|
| **1** | -1.07 ± 0.43 |
| **2** | -1.07 ± 0.49 |
| **3** | -1.13 ± 0.44 |

Table S10: ELS determination of electrophoretic mobility of 102 nm carboxyl-functionalized nanoparticles.

| **102 COOH Replicate** | EPM ($10^{-8}$ $m^2V^{-1}s^{-1}$) |
|---|---|
| **1** | -3.73 ± 0.72 |
| **2** | -4.62 ± 1.35 |
| **3** | -4.43 ± 0.83 |

Table S11: DLS determination of diffusion coefficient of 194 nm unfunctionalized nanoparticles.

| **194 Replicate** | Diffusion Coefficient ($\mu m^2/s$) | Diameter (nm) |
|---|---|---|
| **1** | 2.46 ± 0.78 | 201 ± 82 |
| **2** | 2.47 ± 0.78 | 199 ± 82 |
| **3** | 2.50 ± 0.78 | 197 ± 82 |
| **4** | 2.50 ± 0.78 | 197 ± 82 |
| **5** | 2.55 ± 0.78 | 193 ± 82 |
| **6** | 2.56 ± 0.78 | 193 ± 82 |

Table S12: DLS determination of diffusion coefficient of 198 nm carboxyl functionalized nanoparticles.

| **198 COOH Replicate** | Diffusion Coefficient ($\mu m^2/s$) | Diameter (nm) |
|---|---|---|
| **1** | 2.55 ± 0.78 | 194 ± 82 |
| **2** | 2.58 ± 0.78 | 191 ± 82 |
| **3** | 2.55 ± 0.78 | 193 ± 82 |

Table S13: DLS determination of diffusion coefficient of 196 nm carboxyl functionalized nanoparticles.

| **196 COOH Replicate** | Diffusion Coefficient ($\mu m^2/s$) | Diameter (nm) |
|---|---|---|
| **1** | 2.24 ± 0.52 | 195 ± 43 |
| **2** | 2.22 ± 0.51 | 197 ± 43 |
| **3** | 2.19 ± 0.74 | 199 ± 61 |

Table S14: DLS determination of diffusion coefficient of 102 nm carboxyl functionalized nanoparticles.

| **102 COOH Replicate** | Diffusion Coefficient ($\mu m^2/s$) | Diameter (nm) |
|---|---|---|
| **1** | 4.38 ± 1.53 | 112 ± 54 |
| **2** | 4.68 ± 1.42 | 102 ± 45 |
| **3** | 4.68 ± 1.53 | 104 ± 54 |

Table S15: Surface characteristics of 196 nm COOH-PS NP in NaCl solutions.

| **Ionic Strength (mM)** | **Debye Length (nm)** | **Surface Charge Density ($C/m^2$)** |
|---|---|---|
| **0.25** | 19 | $-4.07 \pm 0.07 \times 10^{-4}$ |
| **0.5** | 14 | $-2.73 \pm 0.04 \times 10^{-4}$ |
| **1.25** | 9 | $-1.46 \pm 0.03 \times 10^{-4}$ |
| **2.5** | 6 | $-1.10 \pm 0.05 \times 10^{-4}$ |
| **5** | 4 | $-0.52 \pm 0.03 \times 10^{-4}$ |

**Table S16:** Characterization of 194 nm unfunctionalized polystyrene nanoparticles in NaCl (0.15-0.35 mM). Mean ± s.d.

| **Ionic Strength (mM)** | **EPM ($10^{-8}$ $m^2V^{-1}s^{-1}$)** | **Diffusion Coefficient ($\mu m^2/s$)** | **Debye length (nm)** | **Charge Number** | **Surface Charge Density ($C/m^2$)** |
|---|---|---|---|---|---|
| **0.15** | -2.3 ± 0.3 | 2.5 ± 1 | 25 | -284 ± 194 | $-3.5 \pm 0.3 \times 10^{-4}$ |
| **0.20** | -2.1 ± 0.3 | 2.6 ± 0.8 | 21 | -222 ± 84 | $-3.5 \pm 0.4 \times 10^{-4}$ |
| **0.25** | -2.1 ± 0.2 | 2.7 ± 0.9 | 19 | -232 ± 126 | $-3.5 \pm 0.3 \times 10^{-4}$ |
| **0.30** | -1.9 ± 0.2 | 2.5 ± 1 | 18 | -212 ± 107 | $-2.8 \pm 0.3 \times 10^{-4}$ |
| **0.35** | -1.5 ± 0.2 | 2.5 ± 0.9 | 16 | -174 ± 80 | $-2.4 \pm 0.3 \times 10^{-4}$ |

Table S17: Surface charge number (Z) values for measured nanoparticles. Mean ± s.d.

| **Particle Type** | **Surface Charge Number** | **Surface Charge Density ($C/m^2$)** |
|---|---|---|
| 194 nm bare | -463 ± 195 | $-6.4 \pm 0.3 \times 10^{-4}$ |
| 198 nm COOH | -503 ± 200 | $-6.6 \pm 0.3 \times 10^{-4}$ |
| 196 nm COOH | -558 ± 208 | $-6.2 \pm 0.4 \times 10^{-4}$ |
| 102 nm COOH | -215 ± 105 | $-1.1 \pm 0.1 \times 10^{-3}$ |
